# Reference-based cell type matching of spatial transcriptomics data

**DOI:** 10.1101/2022.03.28.486139

**Authors:** Yun Zhang, Jeremy A. Miller, Jeongbin Park, Boudewijn P. Lelieveldt, Brian Long, Tamim Abdelaal, Brian D. Aevermann, Tommaso Biancalani, Charles Comiter, Oleh Dzyubachyk, Jeroen Eggermont, Christoffer Mattsson Langseth, Viktor Petukhov, Gabriele Scalia, Eeshit Dhaval Vaishnav, Yilin Zhao, Ed S. Lein, Richard H. Scheuermann

## Abstract

With the advent of multiplex fluorescence *in situ* hybridization (FISH) and *in situ* RNA sequencing technologies, spatial transcriptomics analysis is advancing rapidly. Spatial transcriptomics provides spatial location and pattern information about cells in tissue sections at single cell resolution. Cell type classification of spatially-resolved cells can also be inferred by matching the spatial transcriptomics data to reference single cell RNA-sequencing (scRNA-seq) data with cell types determined by their gene expression profiles. However, robust cell type matching of the spatial cells is challenging due to the intrinsic differences in resolution between the spatial and scRNA-seq data. In this study, we systematically evaluated six computational algorithms for cell type matching across four spatial transcriptomics experimental protocols (MERFISH, smFISH, BaristaSeq, and ExSeq) conducted on the same mouse primary visual cortex (VISp) brain region. We find that while matching results of individual algorithms vary to some degree, they also show agreement to some extent. We present two ensembl meta-analysis strategies to combine the individual matching results and share the consensus matching results in the Cytosplore Viewer (https://viewer.cytosplore.org) for interactive visualization and data exploration. The consensus matching can also guide spot-based spatial data analysis using SSAM, allowing segmentation-free cell type assignment.

## Introduction

Characterizing the spatial distributions of molecularly defined cell types is a shared goal of the Human Cell Atlas, Brain Initiative Cell Census Network (BICCN), and related collaborative efforts. The core elements in this task include transcriptional classification and spatial assignment of cell types, which requires integration of single cell transcriptomics and spatially-resolved transcriptomics to define and match cell type spatially through the analysis of combinatorial gene expression patterns in tissue sections. Single cell RNA sequencing (scRNA-seq) has rapidly progressed into a high throughput standardized methodology and has been used by many labs as a major workhorse for cell type classification in many organs. In contrast, spatial transcriptomics methods are still evolving, varying substantially in methodology, degree of multiplexing, cost, and throughput, lacking consensus data standards and analysis methods.

Characterizing spatially-resolved cell types is essential in the brain in order to study the exceptional cellular heterogeneity and functional significance of its spatial organization. ScRNA-seq has revealed an unprecedented granularity of neuronal cell types in mouse and human brains [1–4], providing a comprehensive landscape of cell type heterogeneity defined by their transcriptional profiles. Recently, a number of multiplex fluorescence *in situ* hybridization (FISH) and *in situ* RNA sequencing methods [5–15] have been reported for conducting spatial transcriptomics experiments at the cellular level. Each method is optimized for marker gene panel design, tissue processing, transcript sequencing, and imaging steps of the pipeline, requiring different strategies for data processing, quality control, and downstream analysis. The SpaceTx Consortium, an organized effort consisting of both experimental and computational working groups, took the lead to evaluate the performance of currently available spatially-resolved transcriptomics methods in high quality cortical samples, with the goal of building consensus maps of cortical cell type distributions based on combined analysis of single cell and spatially-resolved transcriptomics. The overarching effort of the SpaceTx Consortium is summarized in [16].

One aim of the SpaceTx Consortium was to make probabilistic assignments of cell types and map their spatial distributions. Here we describe the quantitative meta-analysis of spatial transcriptomics data with a focus on assigning spatial cell types using the reference cell types from scRNA-seq. This is the first-time spatial transcriptomics data has been analyzed and compared across spatial and computational methods for cell type determination on the same tissue. Here we present the results of these analysis efforts along with strategies for visualization of spatial transcriptomics data. Four available datasets from the SpaceTx Consortium and up to six computational methods are systematically evaluated in the following sections. Available datasets and reproducible work covered in this manuscript are publicly available at the SpaceTx website (https://spacetx-website.github.io/index.html).

## Results

### Analysis overview

We explored multiple approaches to assign the spatial data with reference scRNA-seq cell types, and developed meta-analysis strategies to combine the cell type assignment results from multiple methods to reach consensus assignments (**Figure 1**). We evaluated datasets from four image-based spatial methods (MERFISH [17, 18], smFISH [5, 19], BaristaSeq [20, 21], and ExSeq [15, 22]) in the mouse primary visual cortex brain region (VISp) [1]. All image data (spot-by-gene matrices) were segmented using the same segmentation procedure – Baysor [23], which also included consistent quality control approaches for doublets and low-quality cell removal. The segmentation step produced the cell-by-gene matrices that were used to assign the spatially-resolved cell types to scRNA-seq reference cell types using cell type matching algorithms. Teams of the SpaceTx Consortium explored six computational algorithms (ATLAS [24, 25], FR-Match [26, 27], map.cells* [1], mfishtools [28], pciSeq [29], and Tangram [30]), which produced individual cell type assignments with various probabilistic assignment scores. To arrive at consensus cell type assignments, two meta-analysis strategies were developed to combine the individual assignments more quantitatively (Geometric Mean Combining Strategy, hereinafter GMCS), or more qualitatively (Negative Weighting Combining Strategy, hereinafter NWCS) (**Methods**). In parallel, spot-based cell type assignment was performed by SSAM [31] using a guided mode, which partially borrows information from the combined assignment results. All spatial data and cell type assignment results were loaded into the Cytosplore Viewer (https://viewer.cytosplore.org) for interactive visualization and data exploration, where an integrated tSNE [32] map for all annotated cells in all spatial methods are presented together with single method viewers for comparative analysis.

**Figure 1:**
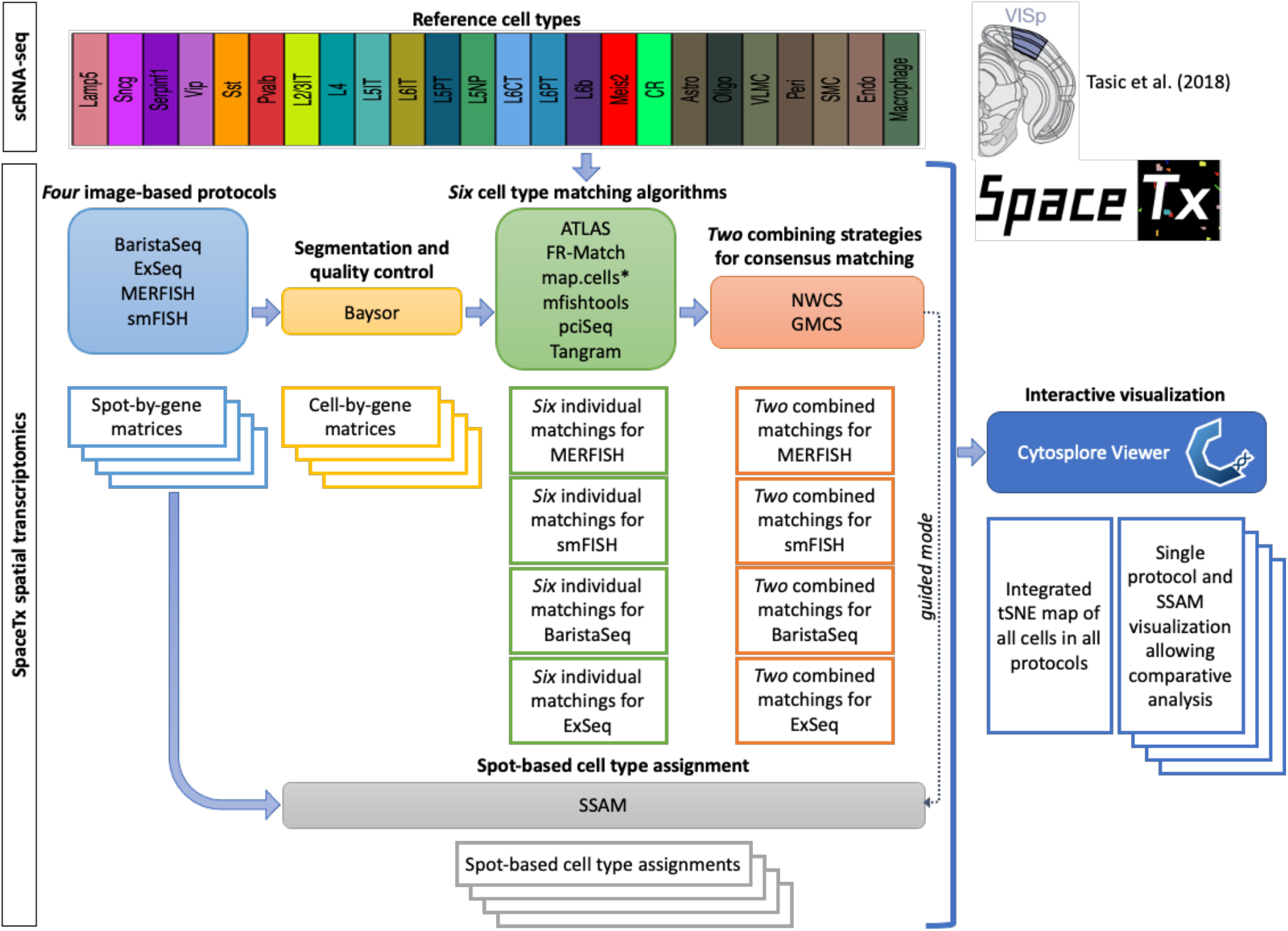
Overview of the SpaceTx analysis workflow. The reference scRNA-seq cell types of the primary visual cortex (VISp) of mouse brain are from Tasic et al. (2018). Spatial transcriptomics data were generated by four image-based experimental protocols (MERFISH, smFISH, BaristaSeq, and ExSeq). Segmentation and quality control were performed using a common procedure (Baysor). Six computational algorithms (ATLAS, FR-Match, map.cells*, mfishtools, pciSeq, and Tangram) for cell type matching were applied. Two meta-analysis strategies were used to combine the individual matching results. Spot-based cell type assignment was conducted using SSAM. All data and matching results can be viewed in Cytosplore Viewer (https://viewer.cytosplore.org).

### Reference cell types

The goal of this study is to produce an initial cell type matching of the spatially-resolved transcriptomics data to open access reference scRNA-seq cell type datasets (a.k.a. scRNA-seq-reference-based cell type assignment of spatial transcriptomics data). The reference mouse visual cortex (VISp) scRNA-seq data were reported in [1], consisting of 14,249 cells with initial 116 cell types defined for VISp. With a focus on spatial gradients, the SpaceTx Consortium re-clustered the data to arrive at a reference cell type taxonomy that contains 191 consensus higher-resolution cell types at the most granular level and 24 cell type subclasses at the intermediate level [16]. With fewer cells per study and fewer reads per cell, the granularity of spatial data collected in this project is not comparable to these most granular scRNA-seq cell types. Therefore, in this study we assigned the spatial data to the cell type subclasses in the reference cell type taxonomy, which distinguishes major GABAergic, glutamatergic, and glial cell types with layer-specific laminar patterning (full list of reference cell type subclasses in **Figure 1**).

### Experimental protocols

As part of the SpaceTx Consortium, tissue sections were successfully collected from mouse VISp and evaluated using MERFISH [17, 18], smFISH [5, 19], BaristaSeq [20, 21], and ExSeq [15, 22] imaging-based experimental protocols. In general, the imaging-based protocols use multi-well plates to stain cells in parallel, and project transcript abundance on microscope images. The spatial methods employ the fluorescent *in situ* hybridization (FISH) technique to localize the transcript sequences. Since each spatial method has unique requirements for numbers of genes and expression levels, each experimental protocol assembles different probe panels with specific gene sets in their design (Supplementary Table S1). The primary output from the imaging-based protocols is a spot-by-gene matrix, quantifying the gene expression intensities in the pixel arrangement of the image.

### Segmentation and quality control

The segmentation step produces a cell-by-gene expression matrix from the image data (spot-by-gene matrix) for downstream analysis. For this purpose, the Baysor algorithm was used because it has been reported to outperform other segmentation tools in terms of yielding better segmentation accuracy, increased number of detected cells, and improved molecular resolution by considering joint likelihood of transcriptional composition and cell morphology [23]. Baysor was applied to perform cell segmentation across all imaging-based protocols to achieve consistent quantification from different protocols. Low quality cells were filtered based on the Baysor cell segmentation statistics, e.g., number of transcripts per cell, elongation characteristics, cell area values, and average confidence scores of segmentation. Cells not passing the quality control filter and cells located outside of VISp based on expert annotation were excluded from further analysis. The final segmented and filtered data (**Table 1**) were used as the input datasets for downstream cell type assignment analysis.

**Table 1:** Summary of segmented and filtered data for each experimental protocol.

| Protocol | Quality control filter | # cells (before / after filtering and annotation) | # genes |
| --- | --- | --- | --- |
| MERFISH | n_transcripts >= 50, elongation < 9, 2000 <= area < 45000, avg_confidence > 0.8 | 6130 / 2150 | 258 |
| smFISH | n_transcripts >= 50, elongation < 8, 500 <= area < 40000, avg_confidence > 0.95 | 4841 / 2360 | 22 |
| BaristaSeq | n_transcripts >= 20, elongation < 8, 10 <= area < 300, avg_confidence > 0.95 | 14095 / 4432 | 79 |
| ExSeq | n_transcripts >= 20, elongation < 10, 5000 <= area < 1500000, avg_confidence > 0.8 | 1504 / 1271 | 42 |

### Comparison of gene properties across experimental protocols

The spatial methods use different reagents, tissue processing steps, barcoding approaches, and amplification methods, resulting in a very different number of genes included in each experiment and different requirements for which genes can be successfully probed (**Table 1**). With these constraints in mind, gene sets were selected to overlap across experiments to the extent possible to allow comparison between studies. We found that while smFISH and ExSeq had relatively fewer genes per experiment (**Table 1**), the average number of transcript molecules detected per cell tended to be higher than for MERFISH or BaristaSeq (**Figure 2A**), even when considering only common genes among experiments (**Figure 2B**).

**Figure 2:**
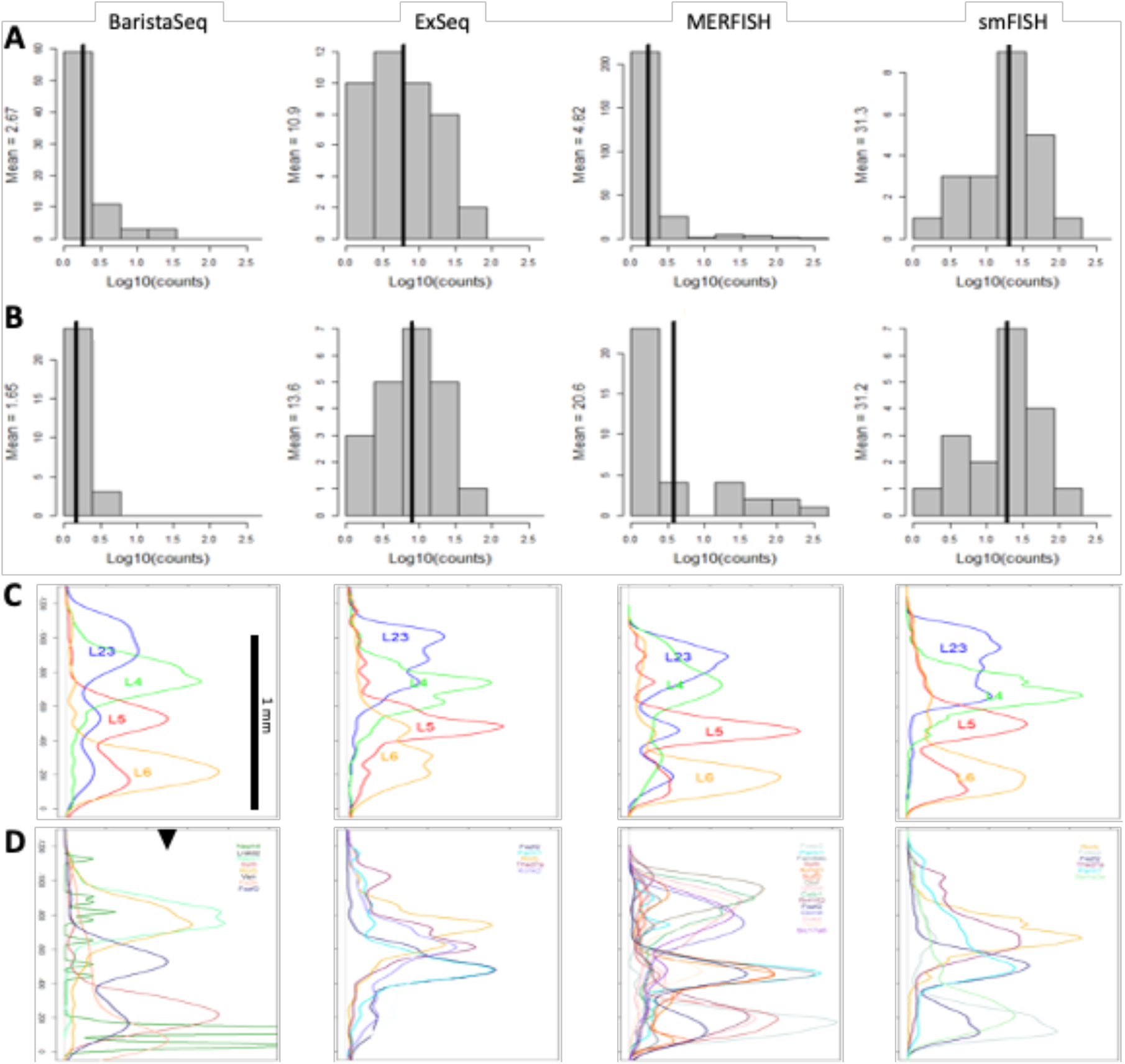
Comparison of gene properties across experimental protocols. (**A**) Distribution of average number of reads in all cells with at least one read for each gene in the experiment. (**B**) Distribution of average number of reads in all cells with at least one read for the subset of genes in the experiment found in at least two other experiments (up to 40 total). (**C**) Density plot of spots across the axis perpendicular to cortical layers (y-axes) for sets of genes marking L2/3 (*Cux2, Lamp5, Cxcl14*), L4 (*Rorb, Rspo1*), L5 (*Fezf2, Parm1*), and L6 (*Sema3e, Foxp2, Syt6*) in mouse VISp. At least one gene from each layer list was assayed in each experiment. Densities (x-axes) are shown in the same scale across all panels in C and D. (**D**) Density plot of up to the 15 genes with the highest maximum density and with maximum density>=0.0025 (black triangle). Genes are color-coded as shown.

Glutamatergic (and to a lesser extent GABAergic) neurons show strong laminar patterning in mouse VISp, and many genes have been well described as showing layer restriction [30], providing a useful ground truth for assessing the accuracy of a subset of genes in each experiment. For all experimental protocols, at least one gene marking L2/3 (*Cux2, Lamp5, Cxcl14*), L4 (*Rorb, Rspo1*), L5 (*Fezf2, Parm1*), and L6 (*Sema3e, Foxp2, Syt6*) in mouse VISp were assayed; in all cases these genes showed maximal expression at the expected cortical depth (**Figure 2C**). Additional computationally-derived genes included in the assays showed layer restriction in MERFISH, and to a lesser extent the other experimental protocols (**Figure 2D**). Together, these results suggest that sufficient information exist from the included gene panels to assign segmented cells to reference cell types at some level of resolution.

### Cell-based cell type matching

Six cell type matching algorithms (ATLAS [24, 25], FR-Match [26, 27], map.cells* [1], mfishtools [28], pciSeq [29], and Tangram [30]) were applied to assign reference scRNA-seq cell types to each segmented cell with an associated confidence score (a.k.a. probabilistic assignment) based on the cell-by-gene count matrix (**Method**). Applying the cell type matching algorithms produces a cell-by-type matching matrix as a primary output, consisting of probabilistic assignment of each segmented cell to each of the reference cell types. For this study, reference cell type subclasses were matched due to the granularity of spatial data limited by the number of marker genes evaluated.

These cell type matching methods each have different advantages and experimental biases, and often produce different cell type assignments, especially in those cells with fewer total transcripts or less confidence segmentation boundaries. To address this, we designed and implemented two metaanalysis combining strategies, each producing a re-calculated confidence score matrix for determining the consensus cell type assignment. The GMCS combined matching considers each individual matching result as the vertex of a polygon whose geometric median, the point with minimum average Euclidean distance from these vertices, serves as the combined result (**Methods**). The NWCS combined matching is a weighted average of the confidence scores from each individual matching method using only the highest score for each cell (**Methods**). For all matching results, deterministic cell type assignment is defined as the cell type with the highest confidence score within a given cell.

#### Individual matchings

In the following two sections, we elaborate the challenges in cell type matching among individual computational methods using the MERFISH data; similarly analysis can be applied to other spatial data as well. A key challenge for the deterministic assignment of cell types was the extensive differences observed among the individual matching results without the availability of a gold standard result to compare against. Confidence scores, as the quantitative metric that reflects the computational matching strength, are however defined differently in each matching method, which showed very different distributional properties (**Figure 3A**). Even though all confidence scores are in the range of [0,1], these scores are not directly comparable across individual matching results because they are very different metrics, for example, correlation or bootstrap probability or p-value, with different distributional properties between the different algorithms. As such, the ranks (i.e., ordered statistics) of the scores are pragmatically more useful, with deterministic cell type assignment using the top-ranked confidence score. The deterministic cell type assignment for the L2/3 IT subclass, for example, further revealed the difference in the number of matched cells (**Figure 3B**) and the spatial distribution of the cells matched to the same subclass in individual matching results (**Figure 3C**). The differences among individual matching results were also reflected in the substantial amount of disagreements of cells matched to the same subclass (**Figure 3D**).

**Figure 3:**
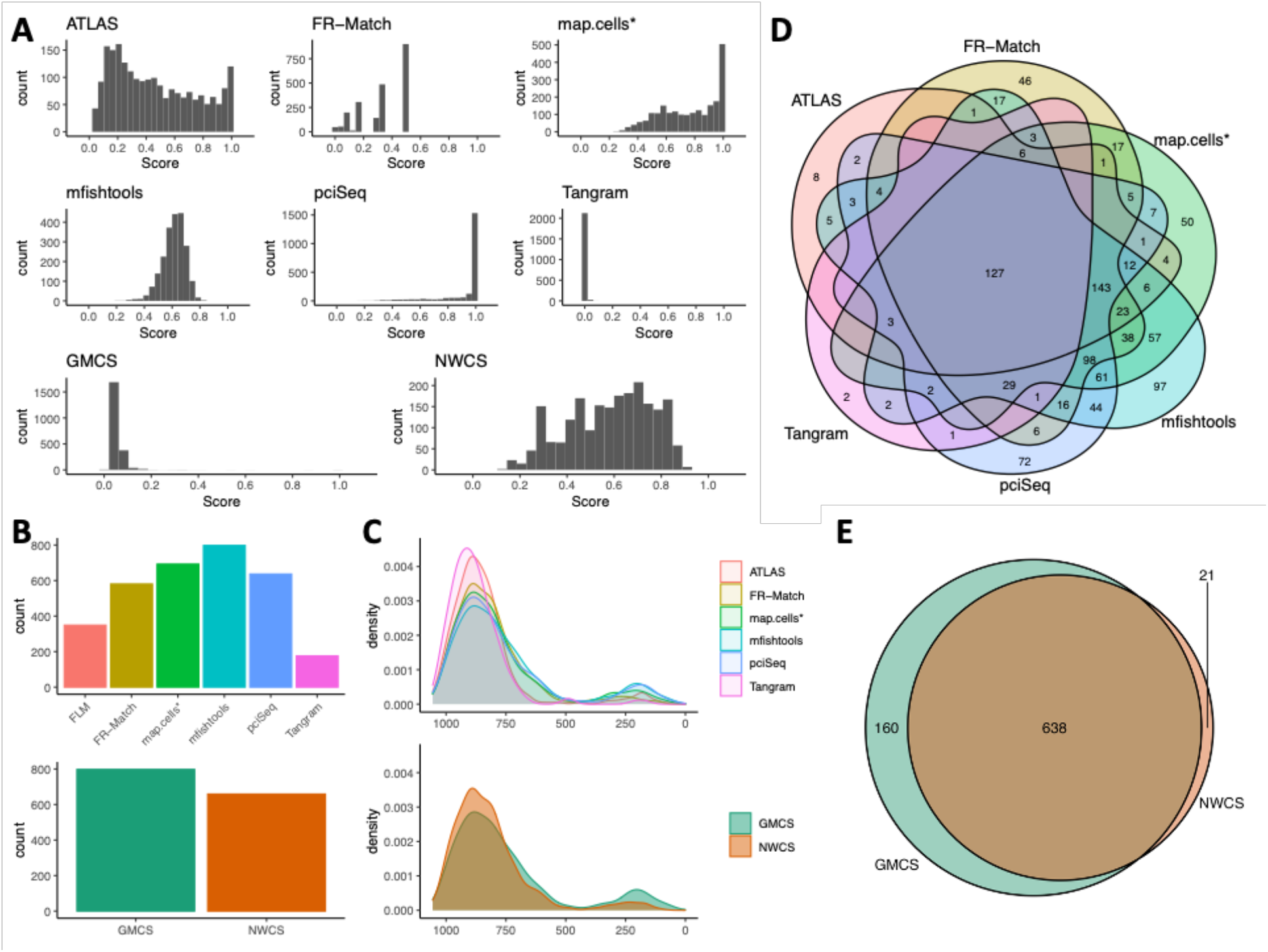
Cell type matching performance comparison on the L2/3 IT subclass of MERFISH data. Six computational methods were applied to match/assign reference cell types to the spatial cells, each resulting in a set of cell-to-type confidence scores ranging from [0,1]. (**A**) The confidence scores from each individual method show very different distributional properties. The ensemble results, GMCS and NWCS, also show very different distributions. (**B**) Number of cells matched to the L2/3 IT subclass by each individual method and ensemble methods. (**C**) Spatial distribution of the cells matched to the L2/3 IT subclass by each individual method and ensemble methods. X-axis is the spatial axis perpendicular to cortical layers (left end: upper layer, right end: deeper layer). (**D**) Overlapping of cells matched to the L2/3 IT subclass by each individual method. (**E**) Overlapping of cells matched to the L2/3 IT subclass by the ensemble methods.

The L2/3 IT subclass is a relatively abundant cell population consisting of intratelencephalic (IT) neurons that are expected to appear in the upper cortical layers 2 to 3. Out of the 2150 MERFISH cells for cell type matching, the number cells matched to the L2/3 IT subclass are 349 (ATLAS), 581 (FR-Match), 693 (map.cells*), 798 (mfishtools), 637 (pciSeq), and 176 (Tangram) in the individual matching results (**Figure 3B**). Though Tangram gives the smallest number of cells matched to the L2/3 IT subclass, most of its matched cells are common cells found in all other matchings, which may suggest that the method for Tangram has high precision (a.k.a. positive predictive value) for this subclass though its detection rate is low (**Figure 3D**). Similarly, ATLAS has the second smallest number of cells matched with high precision. The other four methods matched at least 397 common cells to this cell subclass, suggesting the high precision methods may have a tradeoff of low sensitivity in this case. We may regard the non-common cells specific to each individual method (**Figure 3D**) as the cells that have weaker signal and more noise in their combinatorial marker gene expression pattern; these noisy cells appear to form the major source of the observed spillover effect in the layer distributions (**Figure 3C**) for this specific cell subclass.

That being said, we are not able to conclude that any specific method is better or worse than the others, as all cells are assigned a subclass, among which the noisy cells might have been accidentally assigned to other subclasses but not L2/3 IT subclass in this example. Spatial coordinate plots with confidence score intensities for each individual matching are available in Supplementary Figures S1-S6. All methods were able to recapitulate the laminar pattern of neuronal cells to some extent, but refinements are needed to all of them.

#### Combined matchings

Assuming that the majority of individual methods would produce some level of accurate cell type matching/assignment, combining their results using an ensemble approach may provide the best classification result. We used two different strategies to combine all individual matching results in the ensemble meta-analysis - GMCS and NWCS. Using the L2/3 IT subclass as an example, the combined matchings are more stable in the number of cells (798 for GMCS and 659 for NWCS) matched to the subclass (**Figure 3B**) and more consistent in the spatial distribution of the matched cells (**Figure 3C**). Between the two combined matchings, the vast majority of the cells matched to the same subclass (**Figure 3E**), indicating strong agreement between the two combined matchings. The Combined Matching #1 and #2 assigned 31% and 37% of all MERFISH cells to the L2/3 IT subclass, respectively, though there is still some spillover of the matched cells in the layer distribution. Considering all cells, the two combined matchings produced highly consistent cell type assignment overcoming the large differences among individual matching results, which resulted in 83% (= number of cells assigned the same subclass / total number of cells) of cells being assigned to the same subclass. The combined confidence score intensity matching plots for all cells are available in Supplementary Figures S7-S8; and distribution of all cells in cortical layers by each combined matching are in Supplementary Figure S9. Though the distributions of matched cells in cortical layers are very similar for the abundant GABAergic and glutamatergic subclasses between the two combined matchings (Supplementary Figure S9), they differ in rare and non-neuronal subclasses (e.g., Meis2, Endothelial, and Macrophage), suggesting the increased difficulty for detecting and matching rare cell types in spatial transcriptomics. Overall, these results suggest that, while individual matching algorithms may have different strengths and biases leading to somewhat different results, the ensemble methods via meta-analysis provide a more robust cell type matching/assignment for the spatial cells.

### Spot-based cell type assignment

Working directly on the spot-by-gene matrices, the SSAM [31] framework was used to perform and visualize segmentation-free spatial cell type assignments. Unlike the cell segmentation-based cell type matching/assignment methods, SSAM performs pixel-wise cell assignment which does not require prior cell segmentation, thus independent from the accuracy of cell segmentation. Here, SSAM guided-mode was demonstrated to create cell type assignments, which were guided by the mean log-normalized gene expression of the combined cell type matching results (GMCS and NWCS) (Supplementary Figures S10-S17). In general, the resulting SSAM cell type assignments and the segmented cells with cell type assignments from combined matching results showed visual similarity in both meta-analysis combining strategies for all spatial experimental methods (**Figure 4**). One exception was the GMCS-based cell type assignment of BaristaSeq. This was due to the low quality of the consensus matching; both the segmentation and the SSAM results did not match the previously known layer structure of the visual cortex.

**Figure 4:**
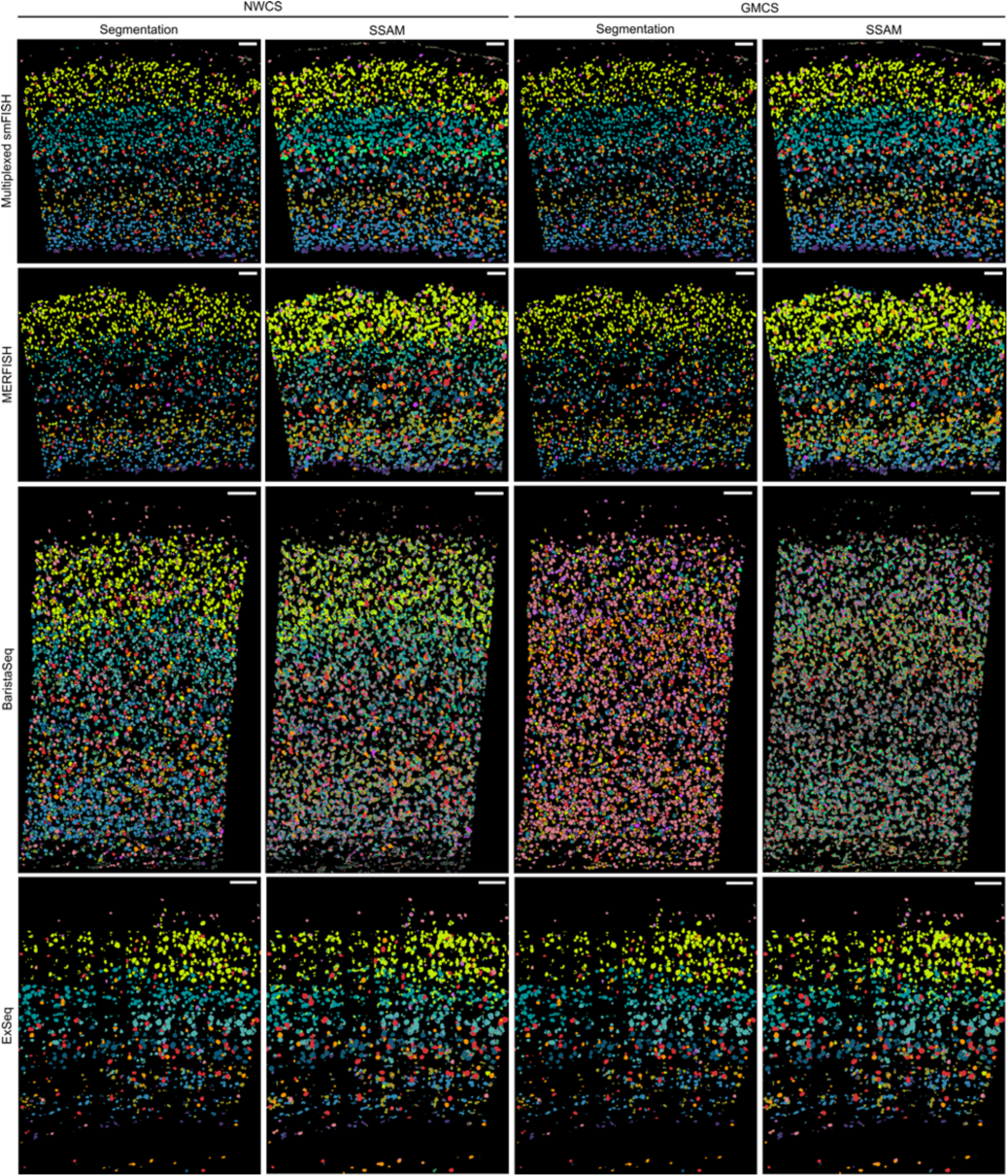
Side-by-side comparison between the segmentation and the SSAM results. The side-by-side comparison between the segmentation and SSAM results show laminar pattern visual similarity in general, which demonstrates SSAM can be used to quickly visualize the spatial cell type distribution without a segmentation step. The colors of each cell type can be found in Figure 1. The scale bars represent 100 μm in all panels.

However, detailed comparison revealed that there are unique spot-based cell type assignments determined by SSAM that were not found in the cell-based approach on the cell segments, e.g., the VLMC subclass in the multiplexed smFISH dataset (olive colored cells in Supplementary Figure S18A). The marker gene of the VLMC subclass *Alcam* showed a very similar gene expression pattern as SSAM identified (Supplementary Figure S18B), which strongly assumes the existence of cells in the region. This observation demonstrates that SSAM could be used as an alternative method to quickly visualize spatial cell types which were possibly missed by prior cell segmentation. Indeed, these results could be used to improve segmentation algorithms. Also, due to the lower density of mRNAs captured by BaristaSeq compared to other FISH methods, the SSAM cell type assignment of BaristaSeq was noisy. Overall, the SSAM analysis shows that a spot-based approach, such as SSAM, is equally capable of revealing good-quality segmentation-free cell type assignment for spatial transcriptomics pixel data, especially using the guided mode analysis when precise gene signatures are given.

Another interesting difference between the segmentation and segmentation-free approach is that, in the smFISH results, SSAM introduced a spatial pattern of a thin layer of spots in deep layer 4 (bright green cells in Supplementary Figure S18C) guided by one segmented cell that was not assigned as L4 in the spatial neighborhood. The segmented cell was assigned to the CR subclass (Supplementary Figure S18A, right panel), a rare and transient class of neurons found in mammalian cortex. Without knowing the ground truth, the assignment of the CR cell could not be validated using the limited number of probe genes in the smFISH panel (22 genes). However, the subtle difference between this cell and the L4 cells were captured from the meta-analysis of the cell type matching results, based on which, SSAM further captured a series of spots show this similar yet distinct expression profile (Supplementary Figure S18D,E). The series of cell spots inferred might be a cell state split from the L4 cells as suggested in [31]. All these observations suggest that segmentation-free approach can provide alternative insights based on the spot data directly.

### Cytosplore Viewer for comparative visualization of spatial protocols

#### ScRNA-seq based gene imputation for data visualization

To compare the different spatial transcriptomics protocols, a combined embedding was generated. A major challenge is that each protocol measures a different set of genes, and the number of shared genes is very small. The four protocols (MERFISH, smFISH, BaristaSeq, and ExSeq) share only six common genes, while the number of genes measured per dataset varies from 22 to 253 genes, where the union of these measured genes contains a set of 314 union genes (Supplementary Table S1). To solve this, we applied SpaGE [33] to impute the expression of the missing genes in each dataset separately and obtain a total of 314 genes per dataset. For each spatial dataset, SpaGE integrates the spatial data with a reference scRNA-seq data measured from the same tissue, and provides prediction for the expression of the missing genes. For example, the MERFISH dataset has 253 measured genes, SpaGE was applied to impute the expression of the 61 remaining genes. Additionally, to reduce batch effects in the joint multi-protocol embedding, we used SpaGE to re-impute the expression of the measured genes in each spatial dataset separately, using a leave-one-gene-out scheme. Taking the MERFISH dataset as an example, SpaGE uses 252 genes for integration with the reference scRNA-seq dataset and provides predicted expression, for the left-out gene, imputed from the scRNA-seq data. This process is applied for all measured genes in each spatial datasets to ensure that all spatial datasets are aligned to the reference scRNA-seq data, and that the expression of all genes is obtained from the same (scRNA-seq) domain. Finally, we generated a combined tSNE embedding for all four spatial datasets using the imputed expression matrix of all 314 genes.

#### Comparative visualization between protocols

For comparative visualization of the different spatial protocols, Cytosplore Viewer (https://viewer.cytosplore.org) was extended with functionality for side-by-side visualization of multiple spatial data sets, in combination with the consensus clustering described above. A comparative view was developed that enables interactive selection from the consensus clustering hierarchy (**Figure 5A**), or from the joint tSNE embedding combining cells from all spatial protocols. Also, either the measured and imputed expression values can be painted on the spatial and tSNE maps enabling comparison of spatial expression patterns across spatial protocols. Finally, functionality for differential expression (DE) analysis between two manual selections (drawn either in the spatial maps or the tSNE maps) was implemented, enabling quick retrieval of differentially expressed genes between regions or cell types. Comparing cell selections within one protocol returns DE of the measured genes for that protocol. Manual selection of cells in the combined tSNE map returns DE of the imputed gene sets.

**Figure 5:**
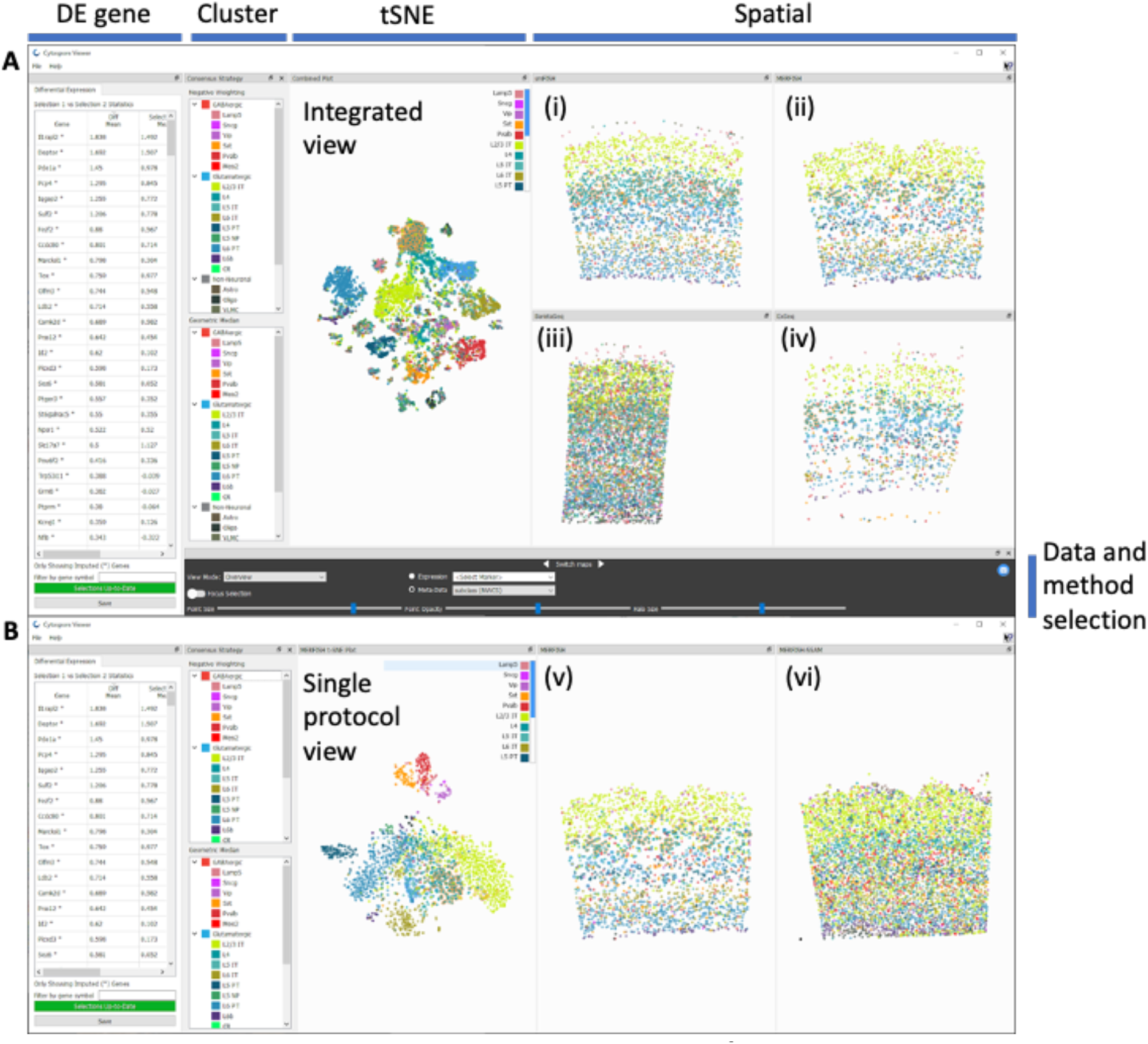
Cytosplore Viewer enables comparative visualization of the SpaceTx data and methods, enabling cell selection from cluster taxonomies (cluster panel), tSNE of single cells based on expression profiles (tSNE panel) and spatial coordinates of cells / local maxima (spatial panels). (**A**) Cross-protocol comparison view: an integrated tSNE map of all cells enables side-by-side comparison of spatial patterning of both consensus matchings on smFish (i), MERFISH (ii), BaristaSeq (iii) and ExSeq (iv), as well as differential expression analysis of cell selections (DE gene panel). (**B**) Single protocol comparison view enables comparing the consensus matchings in the segmentation-based methods (v) and segmentation-free SSAM results (vi) for the individual spatial protocols. Viewing panels are highlighted on the top; data and method selection panel is highlighted to the right of the figure. The NWCS results are shown in both A and B; MERFISH data and results are shown in B. Data and methods can be selected in the data and method selection panel.

#### Single protocol and SSAM visualization

Apart from the comparative visualization, functionality for visualizing individual protocols was developed, consisting of a linked spatial and tSNE map (**Figure 5B**). The tSNE maps were computed on the measured cell-by-gene expression matrices of the individual spatial datasets. Since SSAM omits the step of direct cell segmentation, estimating local correlation between SSAM Kernel Density Estimate profiles and the cluster prototypes, direct comparison between cell-segmented and estimated local correlation maxima was not possible. As such, the individual SSAM local maxima-by-gene matrices were included in the single protocol visualizations.

## Discussion

This manuscript focused on the meta-analysis of cell type matching between spatial transcriptomics data and scRNA-seq reference cell types. The spatial transcriptomics methods are fast evolving, which requires up-to-speed development of data analysis pipelines. Significant emphasis has been devoted to computational algorithms focused on the segmentation step of the imaging-based spatial transcriptomics analysis pipeline [9, 19, 23, 34, 35]; however, limited focus has been on investigating the performance of spatial cell type matching in downstream analyses. This work is the first-time evaluation of scRNA-seq-reference-based cell type matching performance across spatial transcriptomics experimental methods and cell type matching computational methods on the same tissue section.

We first compared gene detection sensitivity and gene expression patterning across spatial experimental methods, which revealed high variability and very different dynamic range in the *in situ* hybridization data across different experimental protocols. We also presented a systematic evaluation of the individual cell type matching algorithms and the combined matching strategies using the MERFISH dataset as an example. The cell-based cell type matching algorithms were applied following the same segmentation step on the image data. Individual matching results varied largely in their metrics of matching confidence as well as their deterministic cell type assignments, among which no overall “winner” could be claimed. Given the variable performance of individual matching results, we used ensemble meta-analysis approaches to combine these individual matchings to form consensus results. The meta-analysis approaches largely improved the agreement between the consensus matchings, where the majority of the cells have the same cell type assignment by the two combined matching strategies. Using the spot-based cell type matching algorithm, similar results as the consensus results could be efficiently obtained without explicit segmentation, given precise gene signatures are available.

A Cytosplore Viewer compilation allows all spatial cells from all evaluated experimental protocols to be viewed in an integrated tSNE map based on the SpaGE-imputed expression scores from scRNA-seq reference data. This enables interactive selection of cells (either through free-form selection or per cell type subclasses), confirming the consistency of the layer patterns across spatial protocols. Differential analysis between free-form cell selections proved particularly useful for identifying gene expression gradients across cortical layers and confirming them across protocols. A side-by-side comparison between the segmentation-based workflow and SSAM revealed a larger density of local maxima detected by SSAM compared to the segmentation-based analysis, however the spatial patterning of cell type subclasses was highly conserved between both methods. Finally, a direct comparison between both combining strategies revealed similar cell type matching results for smFISH, MERFISH and ExSeq. For BaristaSeq, the combined matching by GMCS resulted in inconclusive results, whereas the NWCS matching still performed reasonably well.

The spatial transcriptomics community is growing rapidly with advancements in both experimental and computational methods. For downstream cell type analysis, challenges and opportunities co-exist as well-benchmarked analysis pipelines are lacking. A major goal of the future work would be to promote standardization in data formats and computational methods, including methods for marker selection, probe design, cell segmentation, cell nuclei and boundary delineation, cell type matching, spatial pattern recognition, etc. It will need the community to provide public access to large, high-quality, uniformly collected datasets from all current spatial transcriptomics methods, in common standard file formats, to accelerate innovation in the computational analysis of such data.

## Methods

### Cell-based cell type matching algorithms

We evaluated six computational cell type matching algorithms, namely ATLAS [24, 25], FR-Match [26, 27], map.cells* [1], mfishtools [28], pciSeq [29], and Tangram [30].

ATLAS (A Tool for Learning from Atlas-scale Single-cell multi-omic measurements(ATLAS)) uses a neural network classifier that applies a central moment discrepancy (CMD) [24, 25] term as a domain regularizer to map cell types discover in the scRNA-seq data onto the spatial data. The input to ATLAS is the scRNAseq measurements and the corresponding cell type labels in addition to the spatial transcriptomic measurements at single cell resolution. Using these inputs, ATLAS maps the cell types discovered from the scRNAseq data onto each cell in the spatial transcriptomics data.

The FR-Match algorithm [26, 27] (https://github.com/JCVenterInstitute/FRmatch) requires an initial *de novo* clustering of the spatial transcriptomics data, which provides a supervised mode for the algorithm. Both the candidate spatial cell clusters and the scRNA-seq reference cell types were input to the algorithm, and the best-matched reference cell types for each spatial cell were obtained using the cell-to-cluster function (`FRmatch_cell2cluster`) implemented in the “FR-Match” R package.

The map.cells* algorithm uses a derivative from the map.cells function in “scratch.hicat” R package [1] (https://github.com/AllenInstitute/scrattch.hicat) that was altered to make it more suitable for smaller gene panels. It is a bootstrap-based method that uses Pearson correlation to assess the similarities between cells and cell type clusters.

The mfishtools algorithm [28] (https://github.com/AllenInstitute/mfishtools) also uses Pearson correlation to match cells from spatial transcriptomics method to cell type cluster medians in a scRNAseq reference dataset. This algorithm first applies filtering and scaling strategies to the mFISH and scRNA-seq datasets, and then uses correlation-based assessment to find the best fitting cell type cluster. There are several parameters allowing flexibility in filtering and analysis. Probabilities for cell type assignment were approximated using the following:

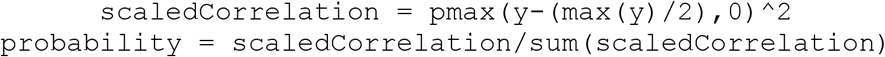

where `y` is the vector of correlations between a given spatial transcriptomics cell and the median expression of each scRNA-seq cell type cluster. Finally, several functions for visualization of matching results and assessment of matching accuracy are included in the mfishtools R package and were applied in this study. A vignette for application of this method is available as part of the “mfishtools” R library.

Probabilistic Cell typing by In situ Sequencing (pciSeq) [29] (https://github.com/acycliq/pciSeq) is a Python package for probabilistic cell typing by *in situ* sequencing. It uses a Bayesian algorithm, leveraging scRNA-seq data to first estimate the probability of each spot belonging to a cell and then each cell to a scRNA-seq cluster. Spots dataframe, segmentation image labels, and scRNA-seq data are required inputs to the algorithm.

Tangram [30] (https://github.com/broadinstitute/Tangram) is distributed as a Python package, based on PyTorch and scanpy. Tangram requires as input a single-cell (or single-nucleus) gene expression dataset and a spatial gene expression dataset. Tangram learns an alignment for the single-cell data onto space by fitting gene expression on the shared genes. The output of the matching algorithm is a cell-by-spot matrix, that gives the probability for cell *i* to be in spot *j*. Using this matching matrix, Tangram can project any annotation (e.g., cell types) from single-cell data onto space. The standard pipeline (with cell-level mapping) has been applied, using functions `tg.map_cells_to_space` for learning the matching and `tg. project_cell_annotations` for projecting cell types computed on scRNA-seq data onto space.

### Combining strategies for consensus matching

#### Geometric Median Combining Strategy (GMCS)

Given the above combining strategy weighing certain matchings over others, we also introduce an independently-developed combining strategy using a geometric median approach that considers each matching equally. Given *m* matchings, each matching *c* cells to a probability distribution over *n* potential cell types, we create a m-gon (polygon with *m* vertices) with vertices in the n-dimensional space (*R^n^*). For each of these polygons, we then find the geometric median, i.e., the point *p* ∈ *R^n^* at which the sum of the *L*_2_ norms from *p* to each vertex in the polygon is minimized. Intuitively, such a point considers each of the individual matchings equally, as having a point *p* closer to one individual matching’s vertex than another would not minimize the sum of the *L*_2_ norms. The confidence with which this matching assigns cell types is consequently a function of how similar or disparate constituent matchings are. Accordingly, certain data modalities for which the individual matching results largely disagree with one another, e.g., BaristaSeq, resulted in not-as-well-classified cells, whereas data modalities in which each cell’s corresponding polygon is of relatively small area, e.g., MERFISH, yielded very well-defined consensus matching (**Results**).

#### Negative Weighting Combining Strategy (NWCS)

A weighting approach was designed to combine the six individual cell type matching results. An evaluation of the individual matching results revealed that: 1) The probabilistic assignments (a.k.a. confidence scores) that reflect the confidence of matching for each spatial cell to each reference cell type showed very different distributions from method to method; some were more binary as either 0 or 1 and others showed more plateau distributions (**Results**). 2) Despite the distributional difference, some cells were assigned to the same cell type with the highest confidence score by all of the methods (i.e., well-matched cells), whereas other cells were only matched to a cell type with a high score by only one method (i.e., poorly-matched cells). In order to avoid the bias introduced by the accidental assignment of those poorly-matched cells, we designed a negative weighting scheme to borrow the best-matched confidence score among all methods. NWCS performs the following steps to combine the individual matching results: 1) Find the best-matched cell types of each cell by keeping the cell-wise highest confidence score. 2) Assign a negative weight (−1) to all other cell types for each cell. 3) The combined confidence score matrix is the sum of all negatively weighted confidence score matrices of each individual method. 4) The NWCS cell type deterministic assignment is the cell type with the maximum confidence score for each cell in the combined matrix.

### Spot-based cell type analysis method

SSAM (Spot-based Spatial cell-type Analysis by Multidimensional mRNA density estimation) [31] analysis is a method that uses the guided mode to generate segmentation-free cell type assignments of the GMCS and NWCS consensus cell types. For all datasets (MERFISH, smFISH, BaristaSeq, and ExSeq), the kernel density estimation (KDE) was performed with the location of mRNAs of each gene with the bandwidth 2.5μm. For SSAM analysis, the resulting vector field was normalized by a library size of 10, and then log-transformed. For GMCS and NWCS cell normalization, the mRNA count of each cell type cluster was normalized to a library size of 10 per cell, and then log-transformed. The gene expression signature of each consensus cell type was computed by taking the mean of all normalized cells in the same cluster. The resulting signatures were then mapped to the vector field, by computing Pearson’s correlations between each consensus signature to all pixels in the vector field. The resulting cell types were filtered with the minimum correlation threshold 0.6.

