## Supplementary Figure for "Reference-based cell type matching of spatial transcriptomics data"

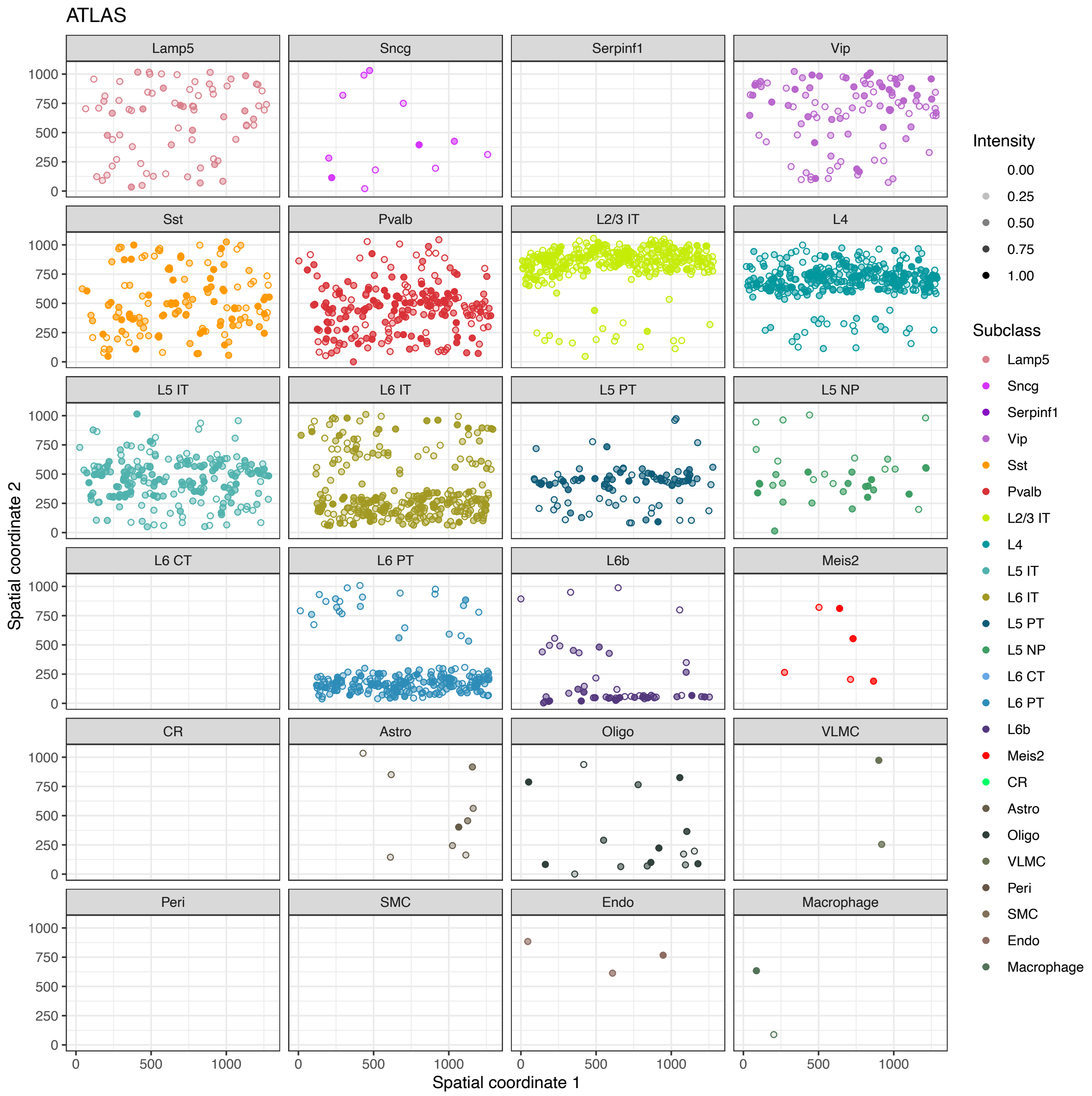


**Supplementary Figure S1:** Spatial coordinate plot by subclass for ATLAS matching results of MERFISH cells.

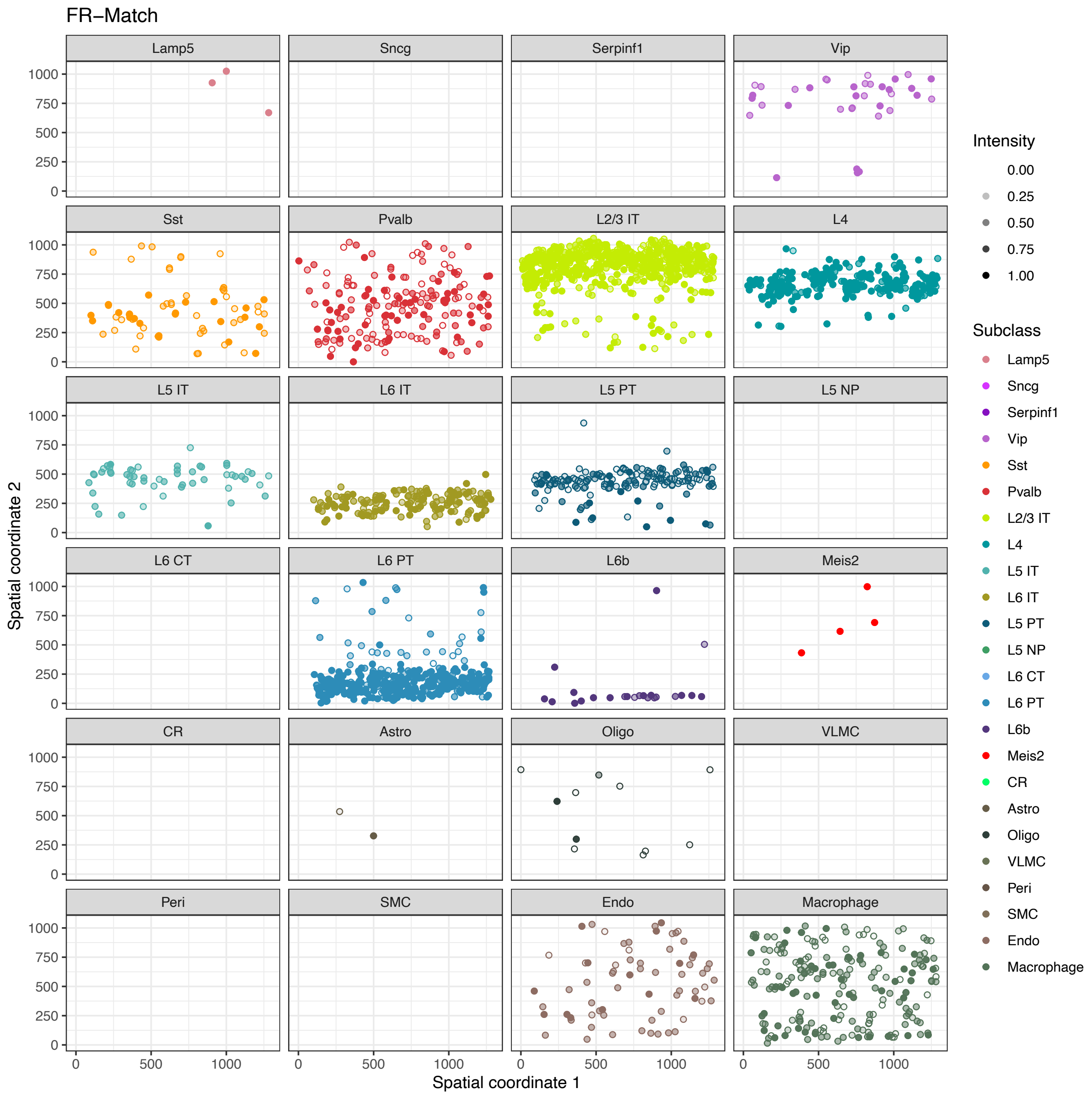


**Supplementary Figure S2:** Spatial coordinate plot by subclass for FR-Match matching results of MERFISH cells.

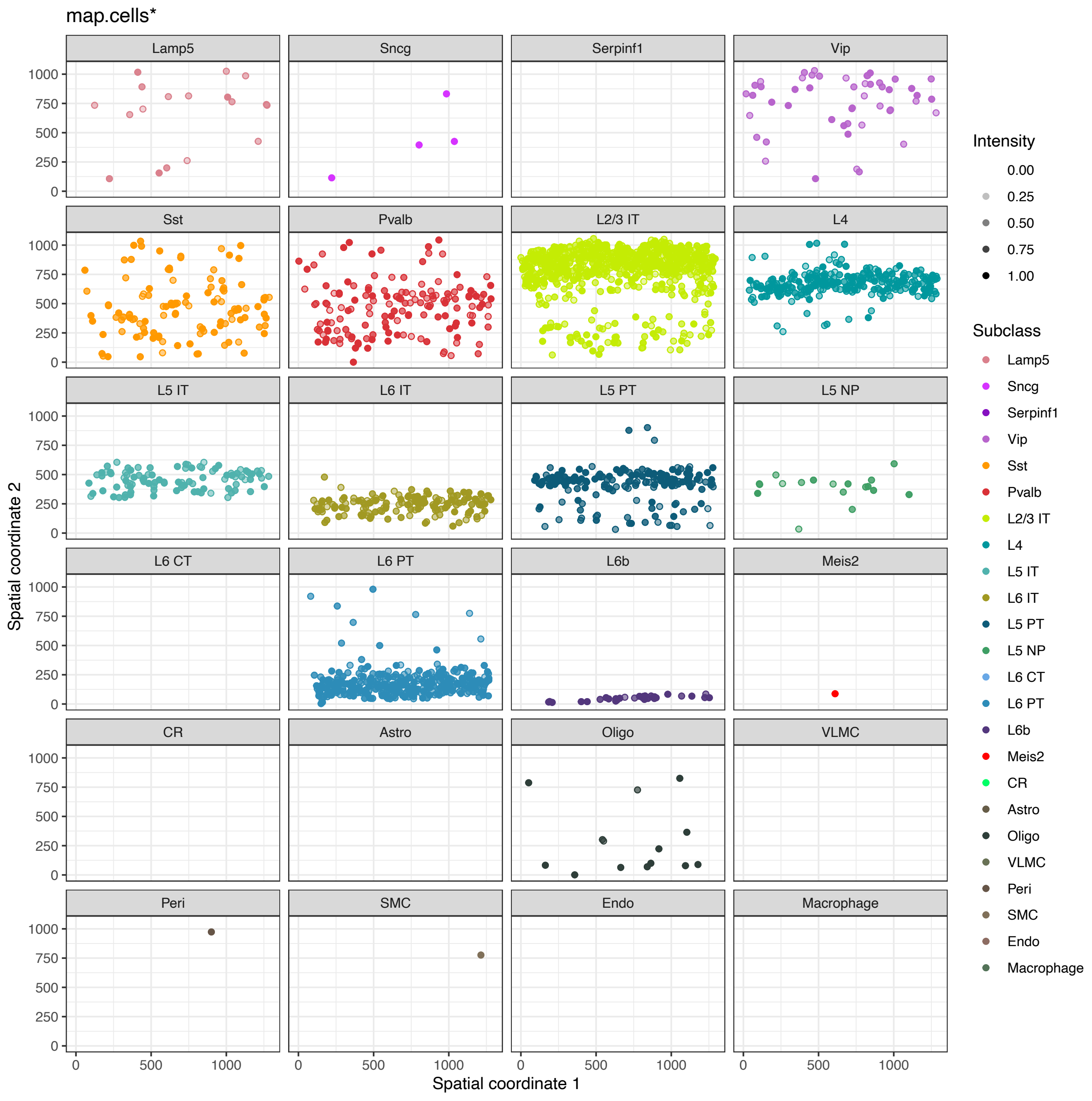


**Supplementary Figure S3:** Spatial coordinate plot by subclass for map.cells* matching results of MERFISH cells.

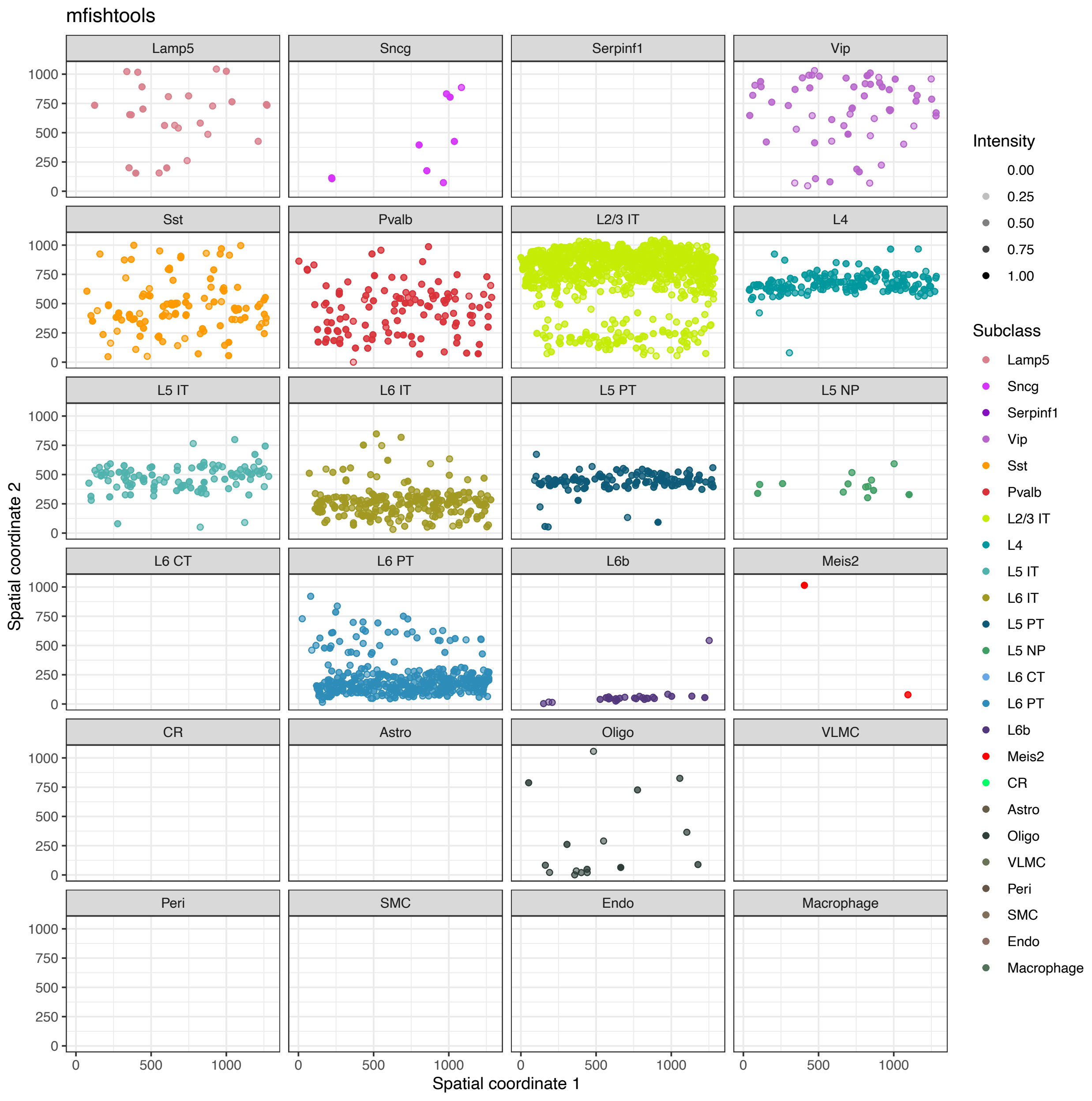


**Supplementary Figure S4:** Spatial coordinate plot by subclass for mfishtools matching results of MERFISH cells.

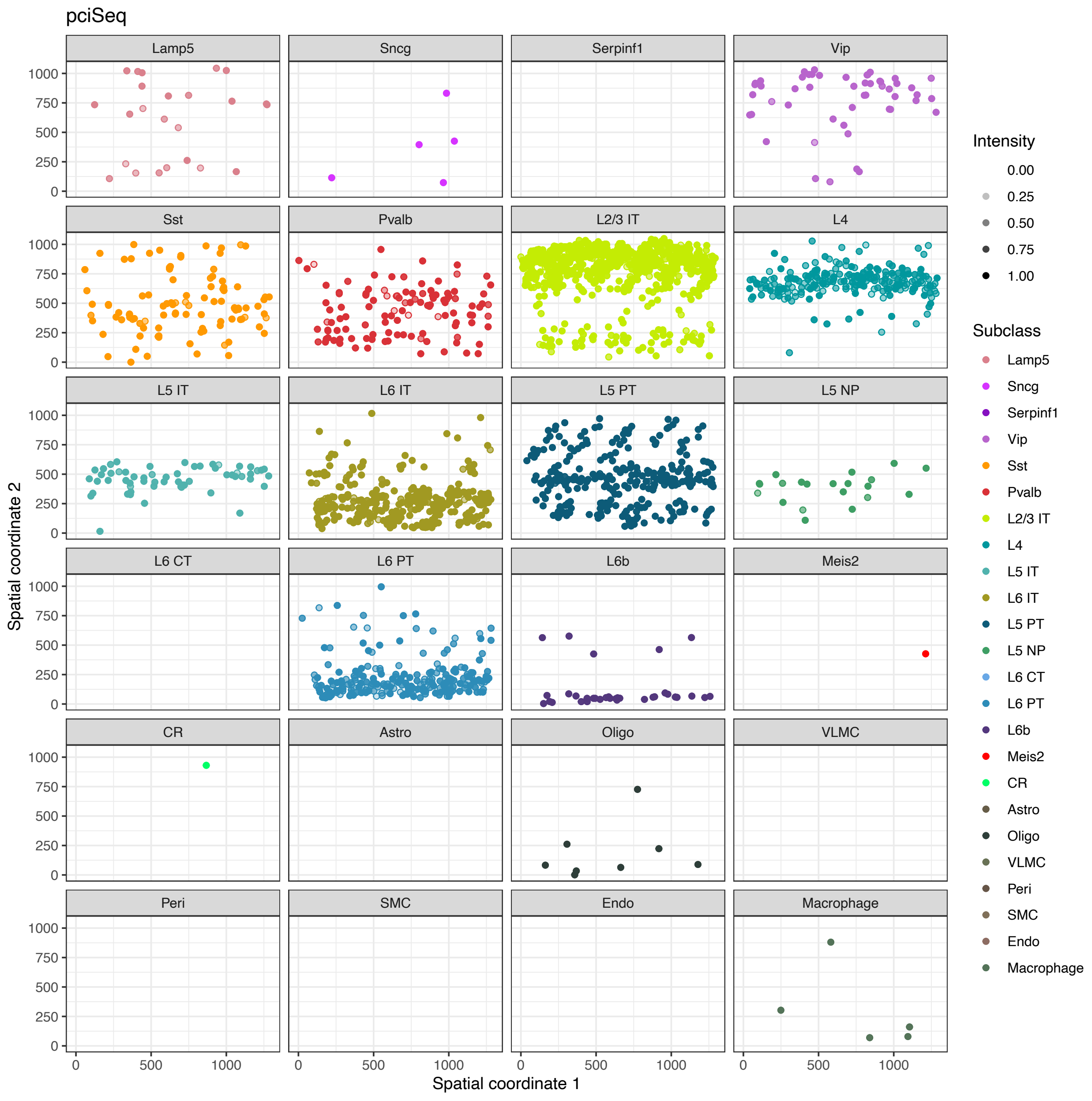


**Supplementary Figure S5:** Spatial coordinate plot by subclass for pciSeq matching results of MERFISH cells.

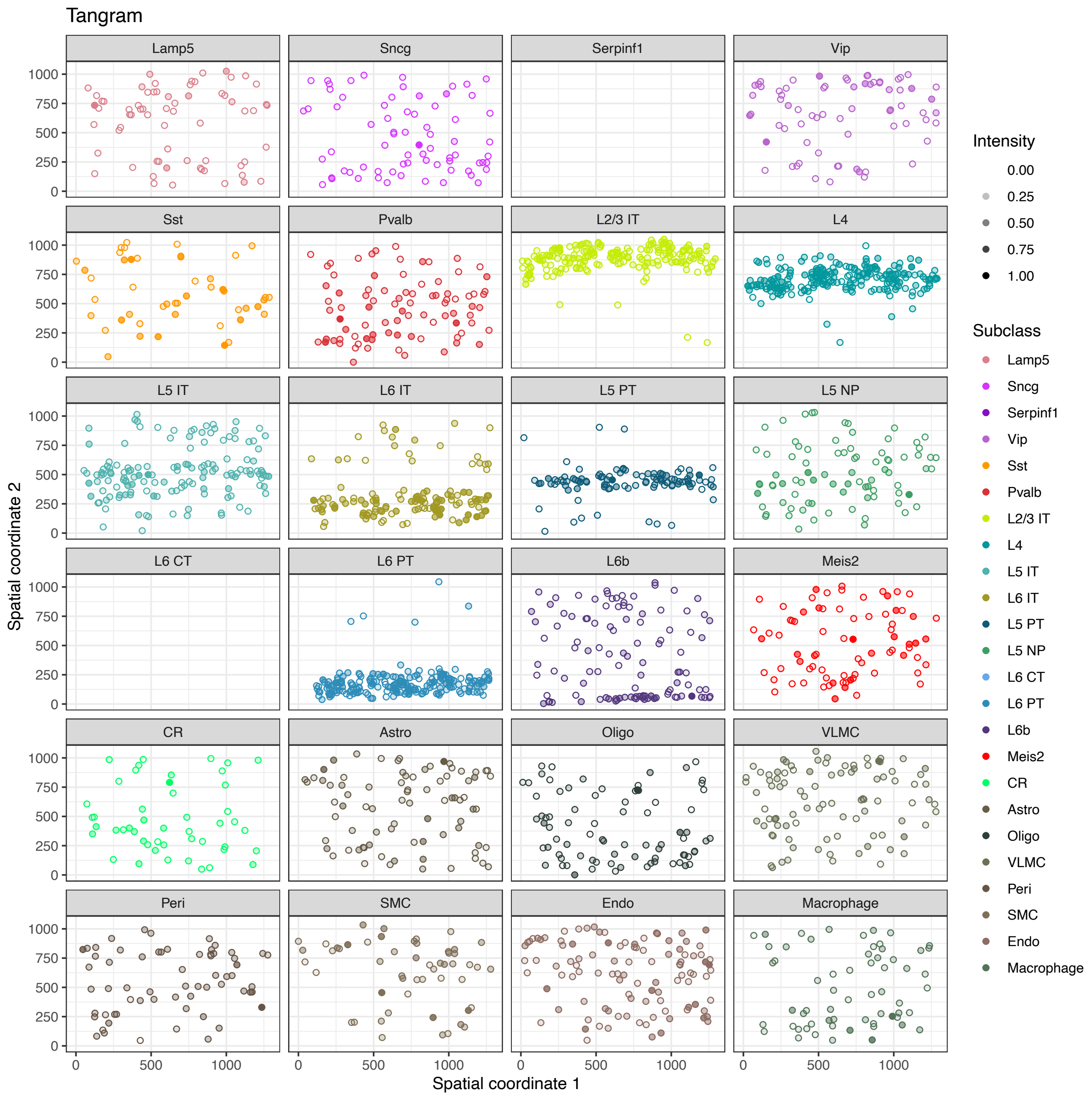


**Supplementary Figure S6:** Spatial coordinate plot by subclass for Tangram matching results of MERFISH cells.

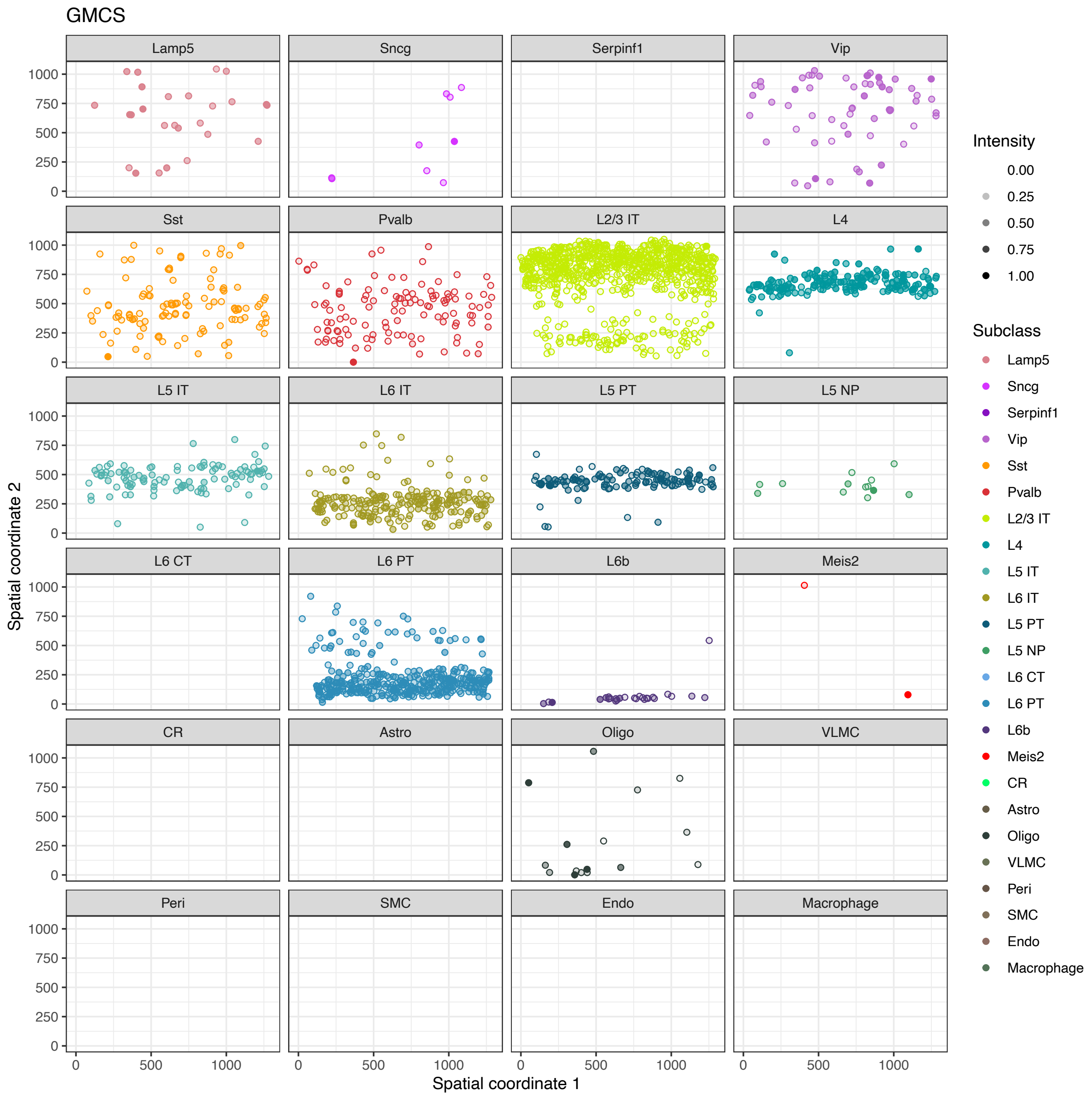


**Supplementary Figure S7:** Spatial coordinate plot by subclass for GMCS matching results of MERFISH cells.

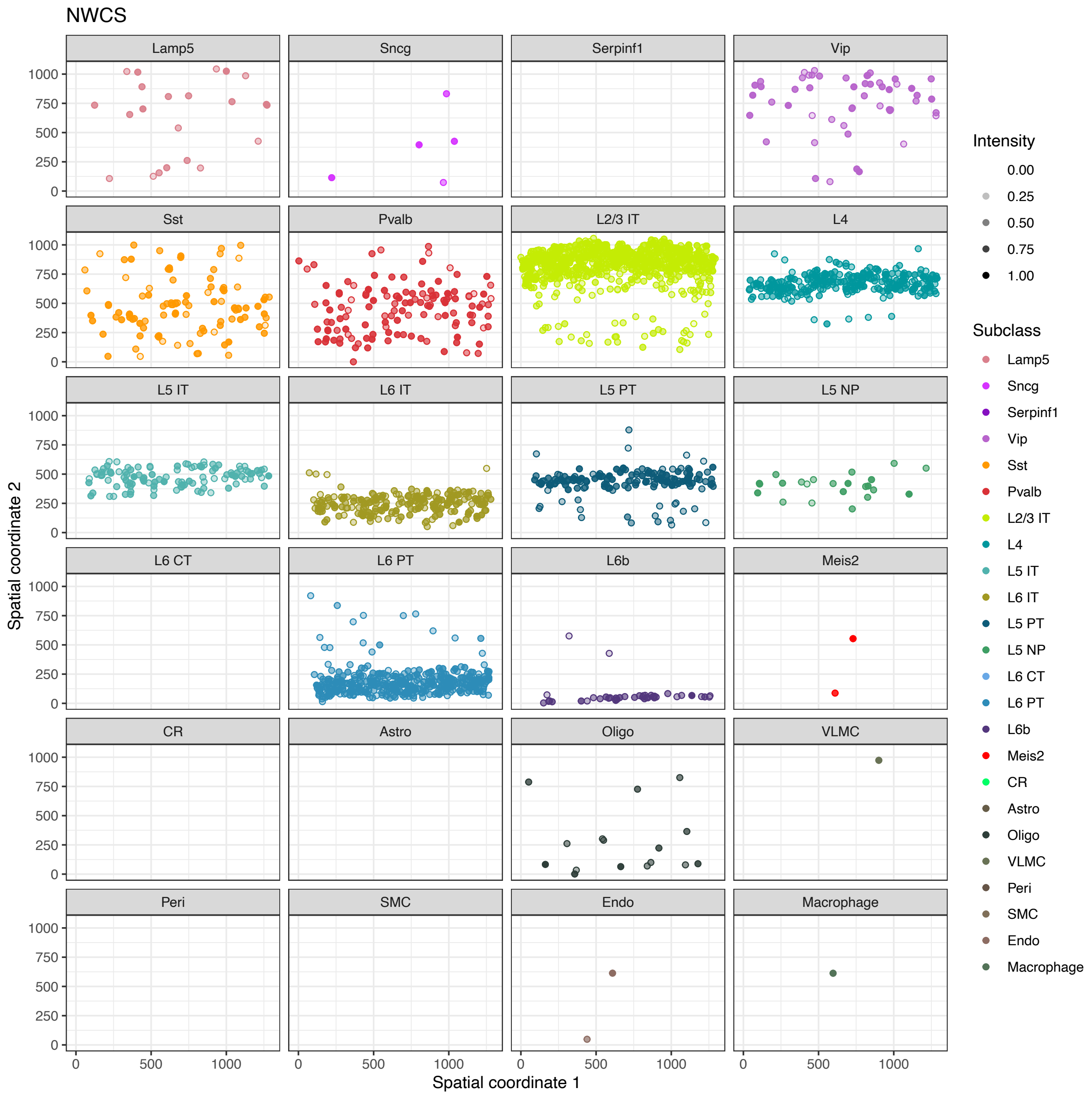


**Supplementary Figure S8:** Spatial coordinate plot by subclass for NWCS matching results of MERFISH cells.

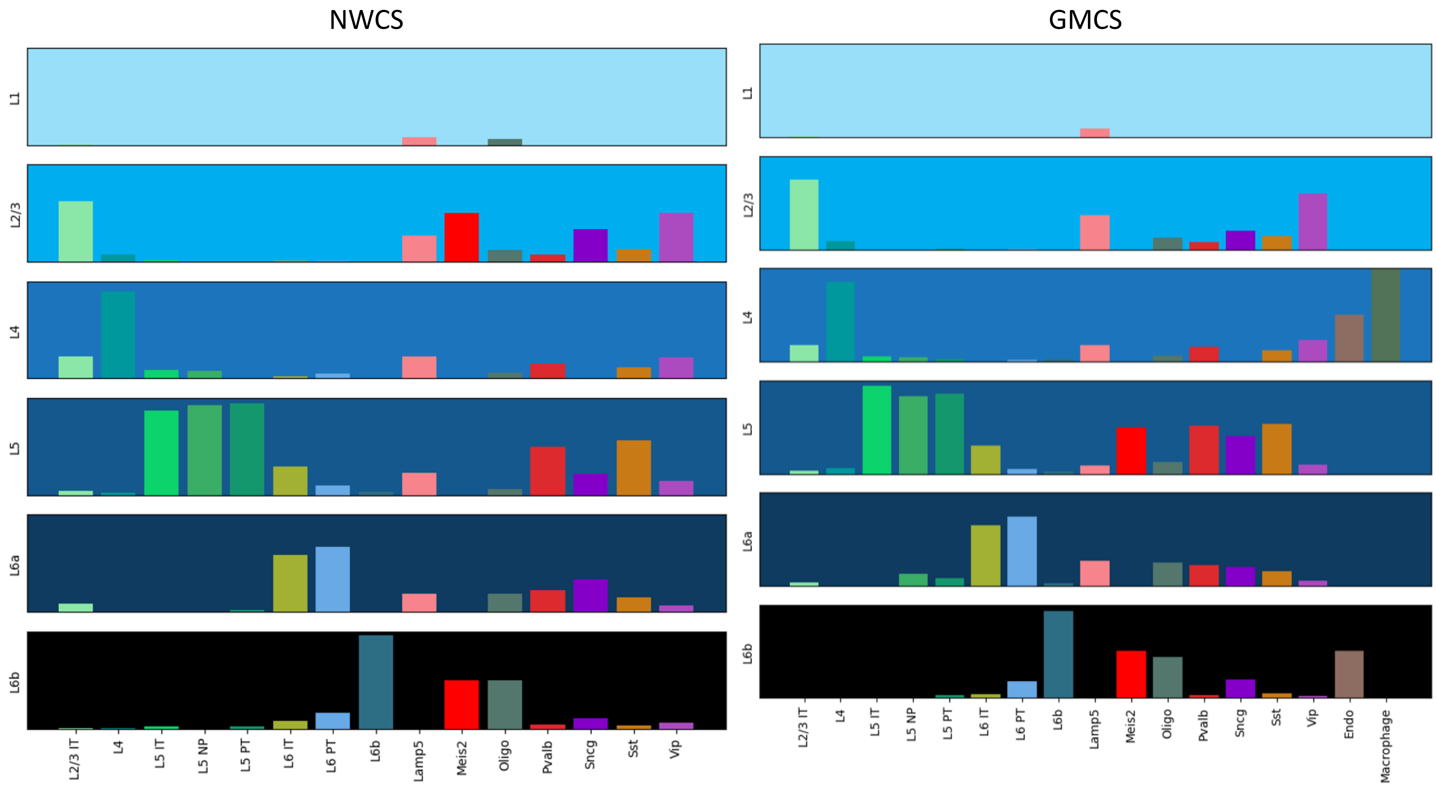


**Supplementary Figure S9:** Cortical laminar distribution of MERFISH cells in the ensembled matching results.

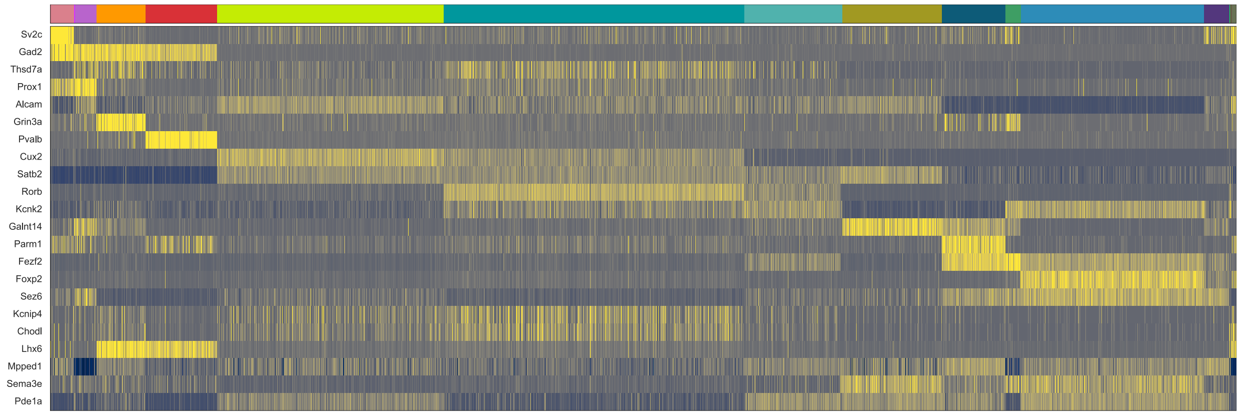


**Supplementary Figure S10:** Heatmap of log-normalized gene expression of smFISH cells clustered using NWCS assigned cell type subclass labels. Rows are smFISH probe genes; column annotation is subclass colors according to Figure 1 in the main manuscript.

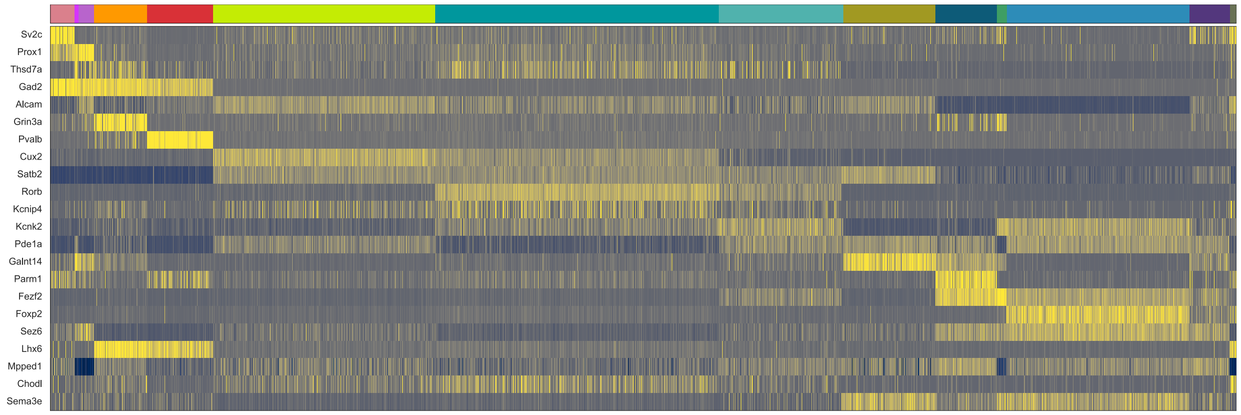


**Supplementary Figure S11:** Heatmap of log-normalized gene expression of smFISH cells clustered using GMCS assigned cell type subclass labels. Rows are smFISH probe genes; column annotation is subclass colors according to Figure 1 in the main manuscript.

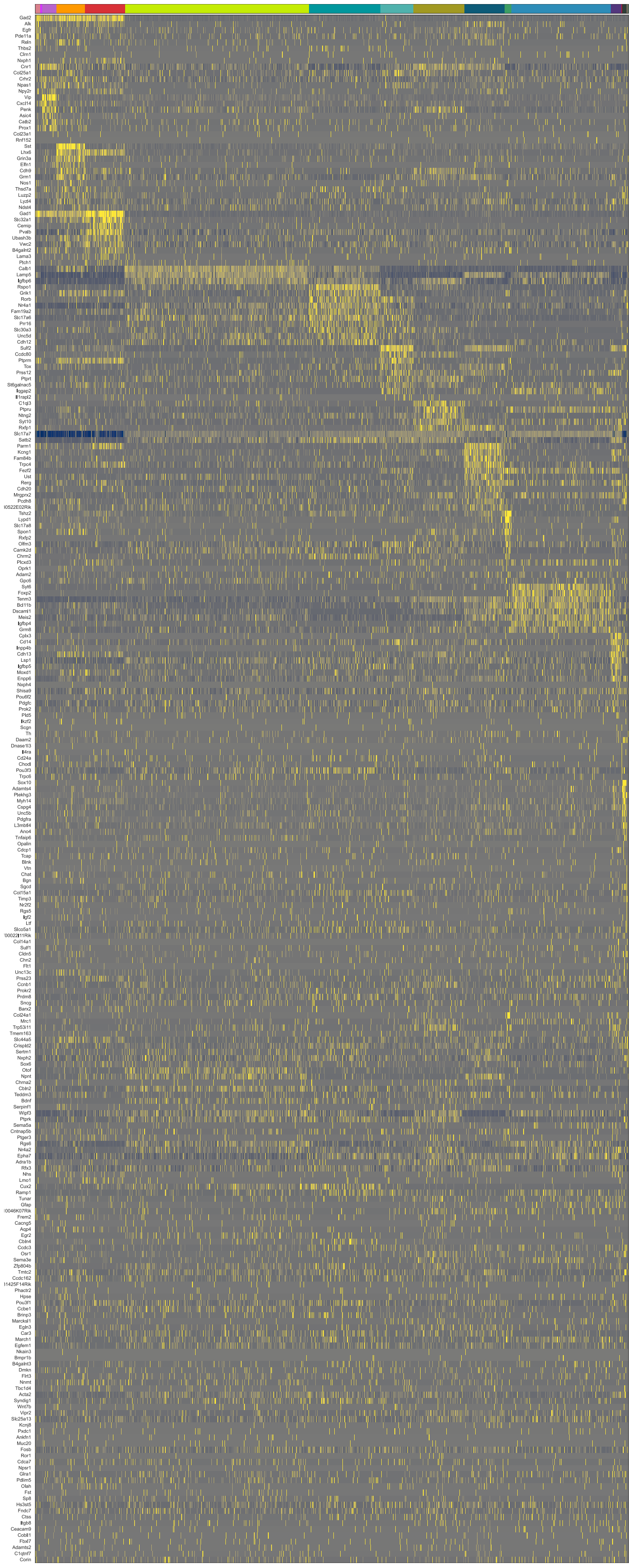

**Supplementary Figure S12:** Heatmap of log-normalized gene expression of MERFISH cells clustered using NWCS assigned cell type subclass labels. Rows are MERFISH probe genes; column annotation is subclass colors according to Figure 1 in the main manuscript.

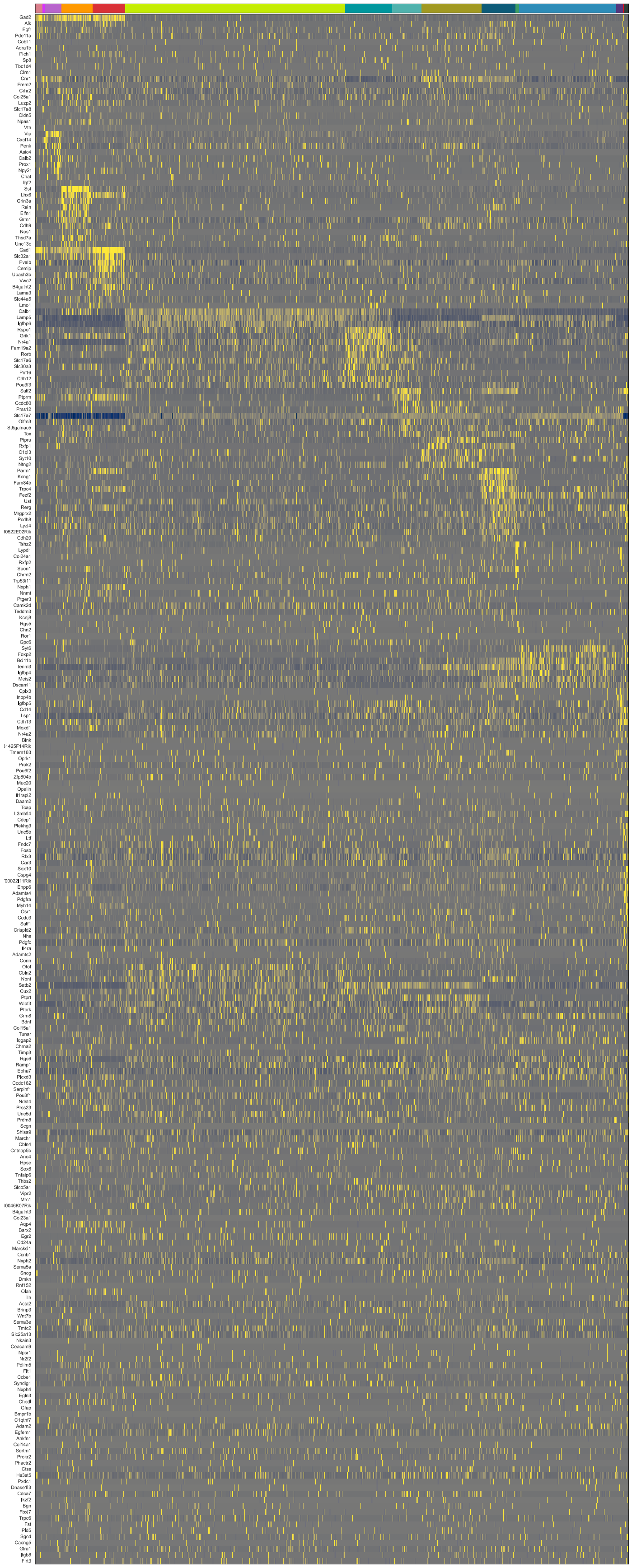

**Supplementary Figure S13:** Heatmap of log-normalized gene expression of MERFISH cells clustered using GMCS assigned cell type subclass labels. Rows are MERFISH probe genes; column annotation is subclass colors according to Figure 1 in the main manuscript.

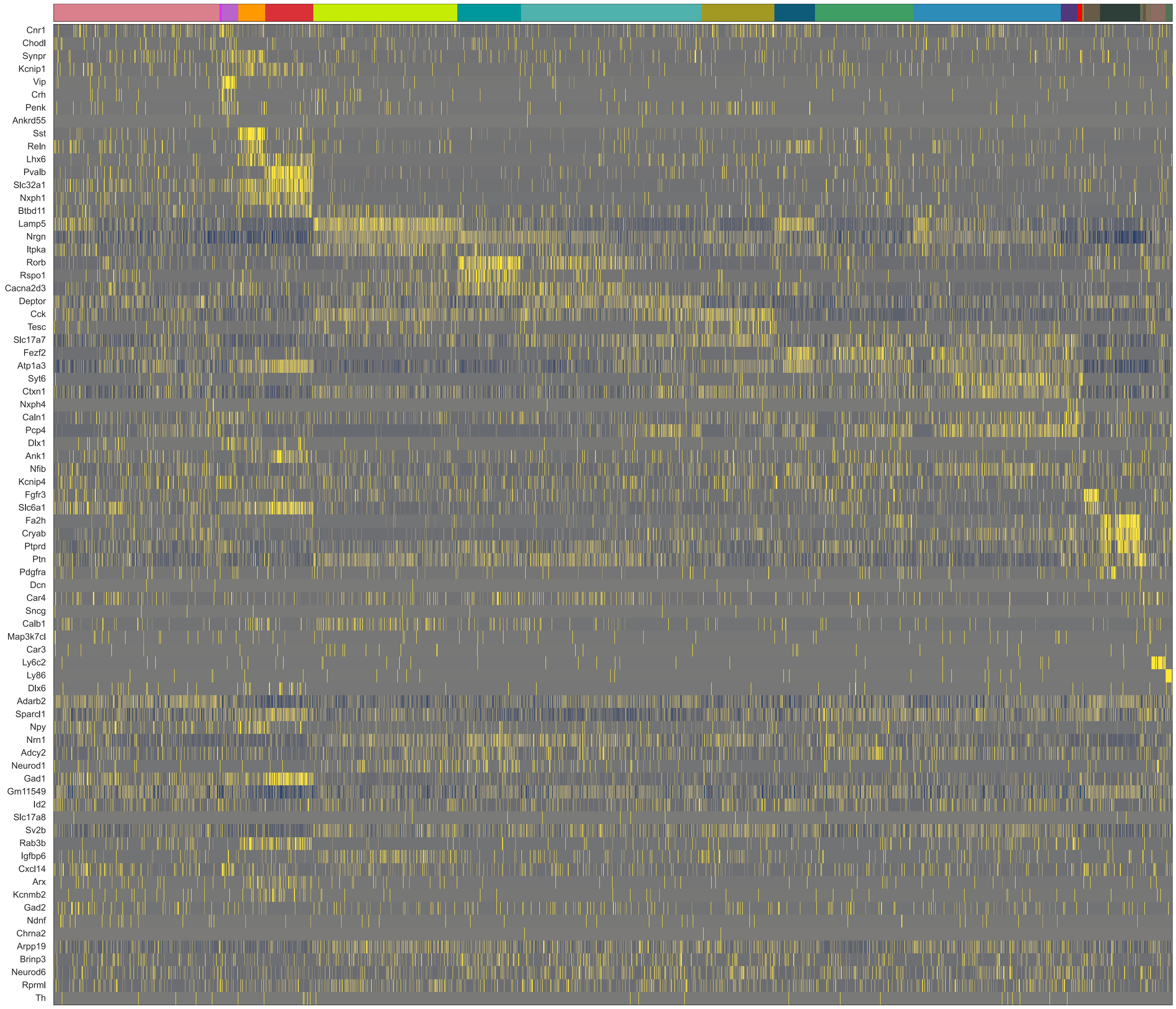


**Supplementary Figure S14:** Heatmap of log-normalized gene expression of BaristaSeq cells clustered using NWCS assigned cell type subclass labels. Rows are BaristaSeq probe genes; column annotation is subclass colors according to Figure 1 in the main manuscript.

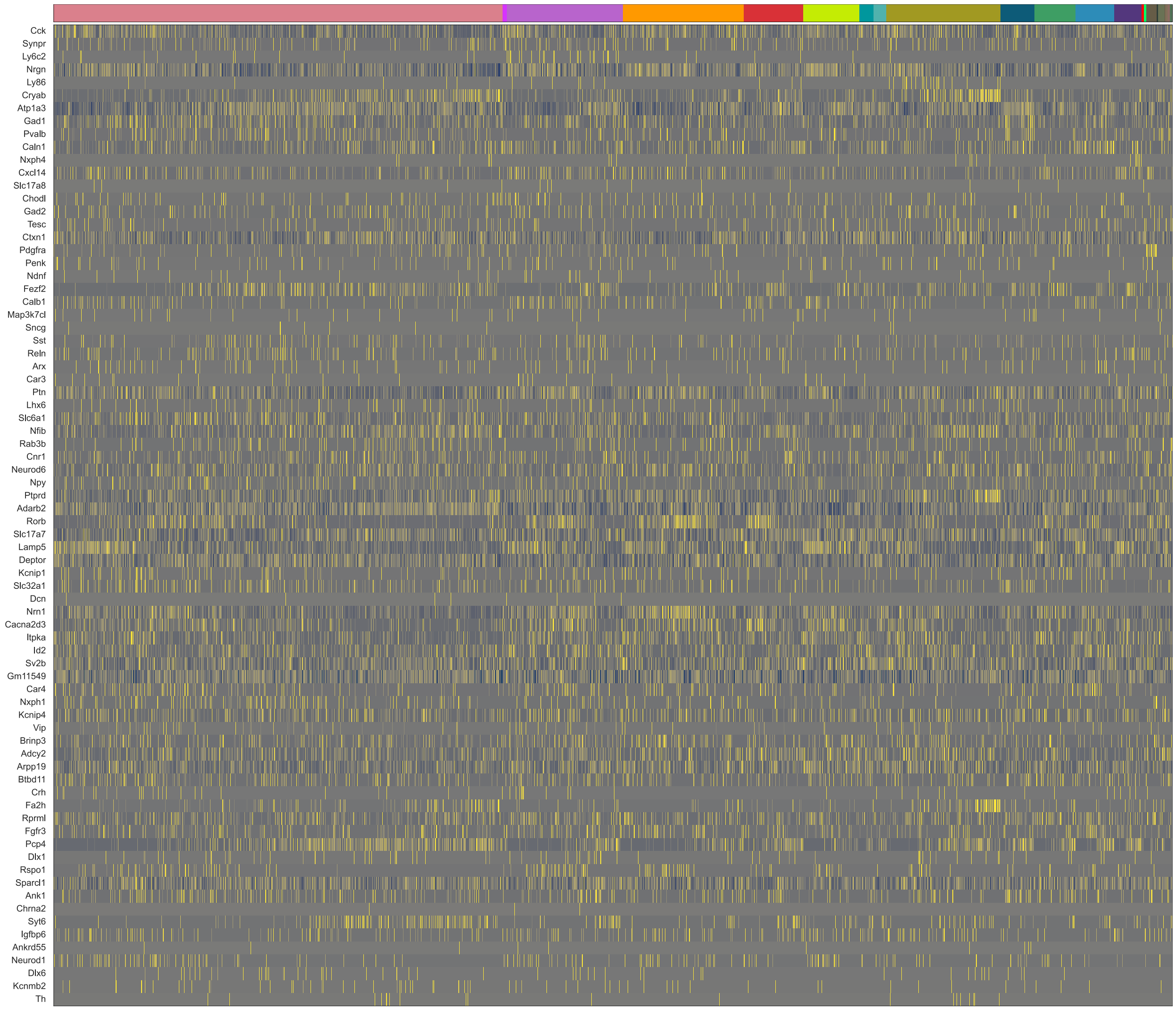


**Supplementary Figure S15:** Heatmap of log-normalized gene expression of BaristaSeq cells clustered using GMCS assigned cell type subclass labels. Rows are BaristaSeq probe genes; column annotation is subclass colors according to Figure 1 in the main manuscript.

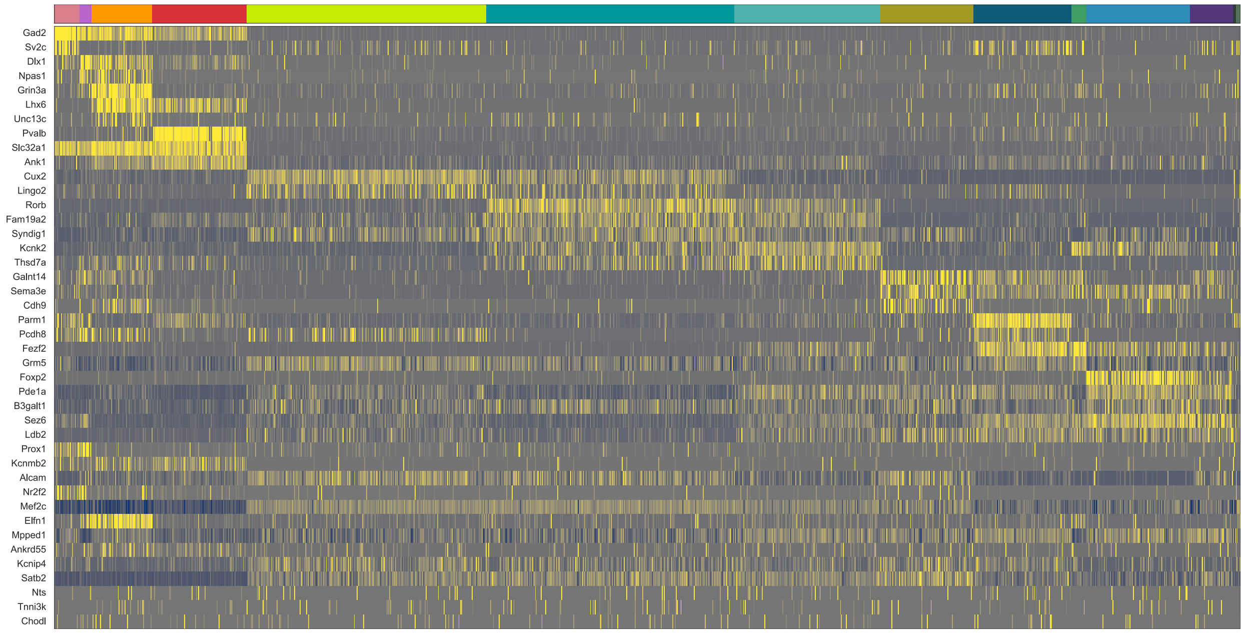


**Supplementary Figure S16:** Heatmap of log-normalized gene expression of ExSeq cells clustered using NWCS assigned cell type subclass labels. Rows are ExSeq probe genes; column annotation is subclass colors according to Figure 1 in the main manuscript.

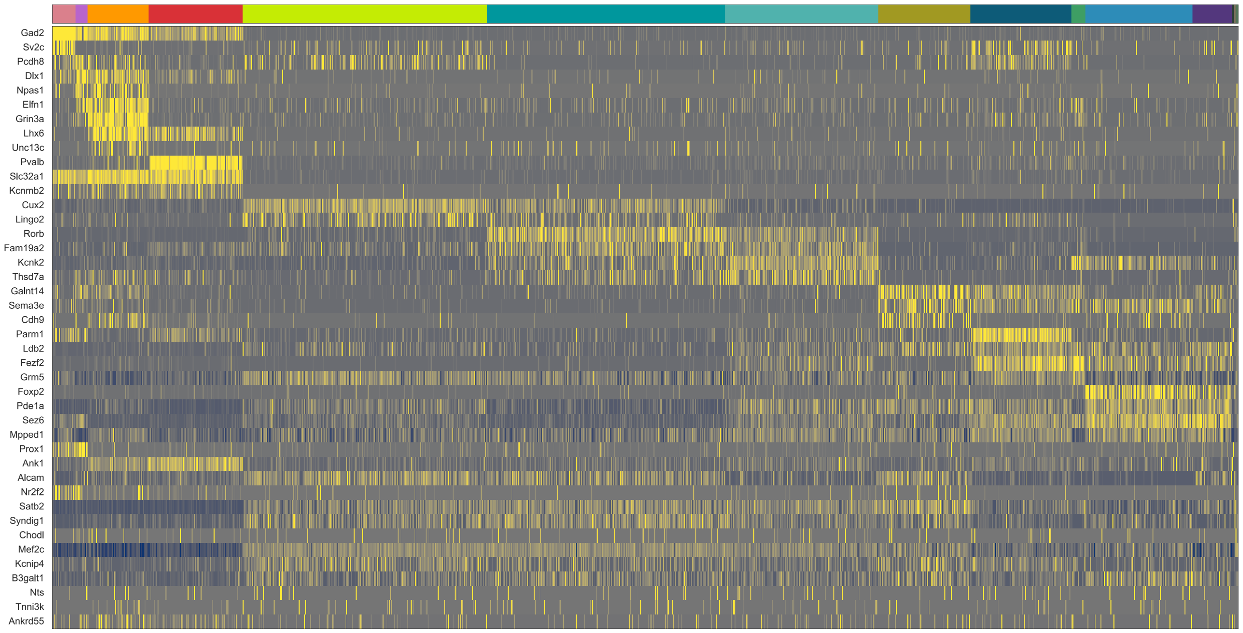


**Supplementary Figure S17:** Heatmap of log-normalized gene expression of ExSeq cells clustered using GMCS assigned cell type subclass labels. Rows are ExSeq probe genes; column annotation is subclass colors according to Figure 1 in the main manuscript.

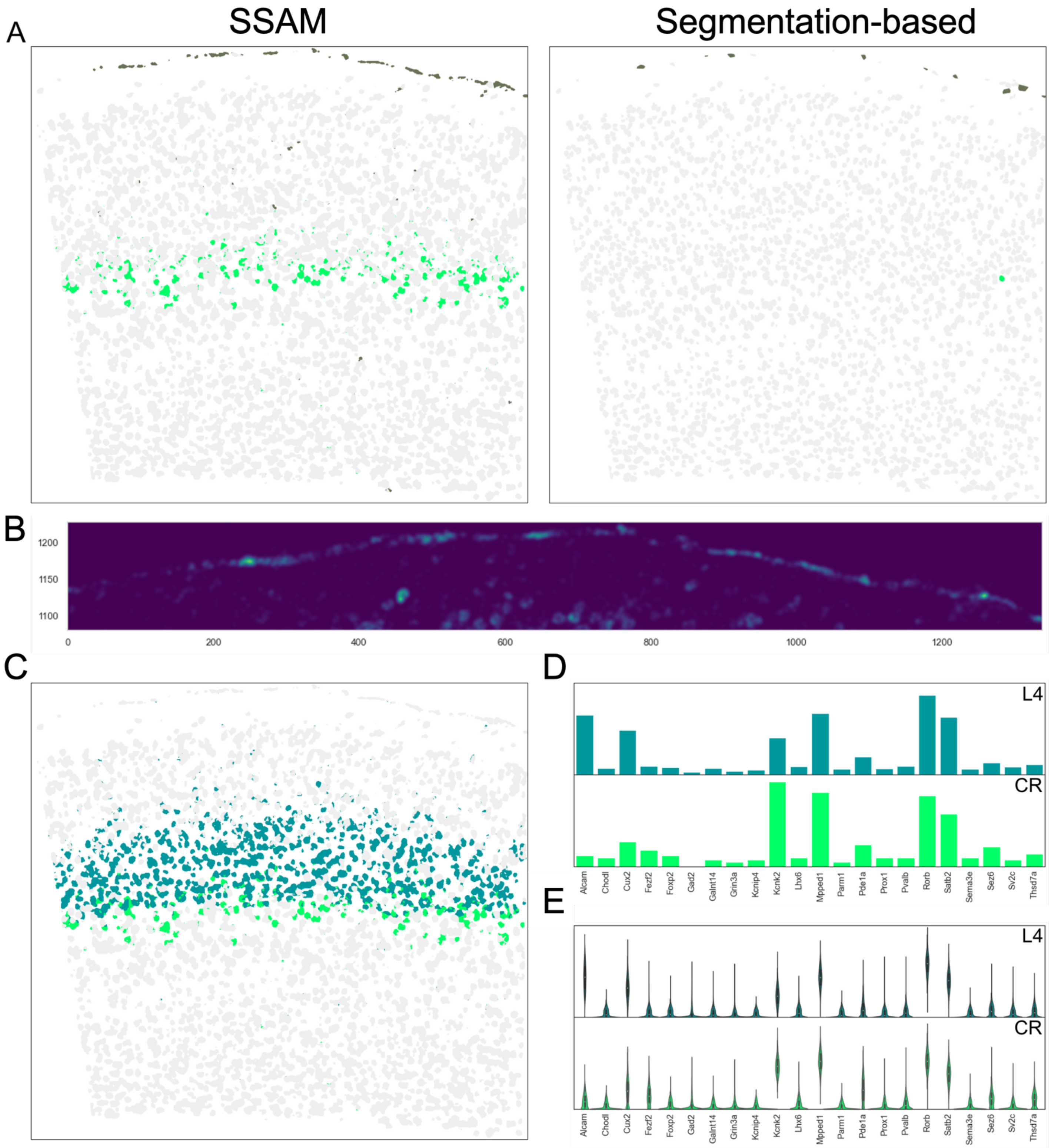


**Supplementary Figure S18:** Detailed comparison between SSAM (segmentation-free) spot-based and segmentation cell-based cell type assignments of the smFISH cells. (A) Side-by-side comparison of VLMC and CR cell types. (B) Gene expression pattern of *Alcam* (a marker gene for VLMC cell type). (C) SSAM cell-type map of L4 and CR cell types. (D) Mean gene expression of cells in the L4 and CR cell types classified by NWCS, as guiding signals inputted to SSAM for spot-based cell type determination. (E) Violin plot of the vectors produced by SSAM for the highlighted cells in panel B.
